# Cellular heterogeneity and lineage restriction during mouse digit tip regeneration at single cell resolution

**DOI:** 10.1101/737023

**Authors:** Gemma L. Johnson, Erick J. Masias, Jessica A. Lehoczky

**Affiliations:** Department of Orthopedic Surgery, Brigham and Women’s Hospital, Boston, MA, USA; Department of Systems Biology, Harvard Medical School, Boston, MA, USA

**Keywords:** blastema, digit tip regeneration, single cell RNAseq, fibroblast heterogeneity, Mest

## Abstract

Innate regeneration following digit tip amputation is one of the few examples of epimorphic regeneration in mammals. Digit tip regeneration is mediated by the blastema, the same structure invoked during limb regeneration in some lower vertebrates. By genetic lineage analyses in mice, the digit tip blastema has been defined as a population of heterogeneous, lineage restricted progenitor cells. These previous studies, however, do not comprehensively evaluate blastema heterogeneity or address lineage restriction of closely related cell types. In this report we present single cell RNA sequencing of over 38,000 cells from mouse digit tip blastemas and unamputated control digit tips and generate an atlas of the cell types participating in digit tip regeneration. We define the differentiation trajectories of vascular, monocytic, and fibroblastic lineages over regeneration, and while our data confirm broad lineage restriction of progenitors, our analysis reveals an early blastema fibroblast population expressing a novel regeneration-specific gene, *Mest*.

## INTRODUCTION

Many animals have the amazing ability to regenerate entire body parts such as the limb, tail, or spinal cord following amputation. This process has been termed epimorphic regeneration, where a complex structure comprised of multiple tissue types is regenerated from progenitor cells within a structure termed the blastema (Carlson, 1978; Hay and Fischman, 1961; Morgan, 1901). Examples of vertebrates that employ epimorphic regeneration include axolotl, newt, and juvenile xenopus which can regenerate many structures including limbs and the spinal cord (Dent, 1962; Overton, 1963; Spallanzani, 1768); and zebrafish, which can regenerate their fins (Johnson and Weston, 1995). In contrast to these species, mammals have limited epimorphic regeneration of complex tissues, though examples do exist: deer can repeatedly shed and regenerate antlers, and mice and human children can regenerate amputated digit tips (Goss, 1961; Illingworth, 1974; Neufeld and Zhao, 1995). Mouse is a well characterized model for studying digit tip regeneration. Following amputation in adult digit tips, there is an initial inflammation and wound healing phase (Fernando et al., 2012). When the wound epithelium has closed, the blastema, a proliferative and heterogeneous structure, forms and goes on to regenerate all non-epidermal structures of the digit tip by approximately 28 days post-amputation (dpa) (Fernando et al., 2012; Lehoczky et al., 2011; Rinkevich et al., 2011).

The blastema is the common structure that links together regeneration in species that seem disparate such as zebrafish, axolotl, and mouse. The blastema is a critical yet transient structure and much remains to be learned about how it mediates regeneration of complex tissues, particularly in mammals. Two hypotheses exist as to how blastema cells give rise to regenerated tissues. One posits that blastema cells are multipotent and can differentiate into any of the regenerating tissues. Another is that the blastema is a heterogeneous population of cells that are lineage restricted, and only contribute to their tissue of origin in the regenerate. While species-specific nuances likely exist, genetic lineage tracing studies in several regenerative models support that the blastema contains cells that are lineage restricted, not multipotent (Flowers et al., 2017; Gargioli and Slack, 2004; Tu and Johnson, 2011). In the mammalian digit tip specifically, mouse genetic lineage analyses have revealed that embryonic germ layer identities hold true during digit tip regeneration and there is no evidence for transdifferentiation (Lehoczky et al., 2011; Rinkevich et al., 2011). Progeny of epithelial progenitor cells traced using *Krt5* or *Krt14* inducible cre drivers remain restricted to the regenerated epithelium (Lehoczky et al., 2011; Rinkevich et al., 2011; Takeo et al., 2013). Similarly, *Sp7* or *Sox9* marked osteoprogenitors contribute solely to the regenerating bone and periosteum, and *VE-cadherin-* or *Tie2-*expressing endothelial cells only give rise to endothelium in the regenerate (Lehoczky et al., 2011; Rinkevich et al., 2011). In the neural lineage, Schwann cells marked by *Sox2* contribute only to the regenerated glial lineage (Johnston et al., 2016). Fibroblasts are one of the most abundant cell types within the digit tip, and as found for the other cell types, lineage marked *Prrx1*-expressing fibroblasts remain fate restricted to the regenerated mesoderm (Rinkevich et al., 2011). In a similar experiment, *Msx1*-expressing cells in the mesenchyme and bone contribute highly to the blastema but do not transdifferentiate into tissues lineages derived from other germ layers (Lehoczky et al., 2011). Collectively, these studies support a heterogeneous blastema comprised of lineage restricted progenitor cells in the regenerating mouse digit tip. However, these previous analyses lack a precise description of all the cell types present in the blastema and an assessment of lineage restriction among closely related cell types.

To this point, previous genetic lineage analyses in axolotl found the regenerating limb blastema to be heterogenous and lineage restricted (Kragl et al., 2009), and recently single cell RNAseq and lineage tracing have been combined to elucidate a more detailed understanding of the axolotl limb blastema (Gerber et al., 2018; Leigh et al., 2018). For example, the presence of macrophages, muscle progenitors, and fibroblasts was confirmed while additional cell types were discovered in both regenerating and homeostatic limbs (Leigh et al., 2018). Moreover, supporting transdifferentiation of closely related lineages, a multipotent fibroblast-like progenitor competent to contribute to multiple regenerated lineages including tendon, skeleton, and fibroblasts was found in the blastema (Gerber et al., 2018). However other lineages, including muscle and wound epithelium, remained more restricted (Gerber et al., 2018; Leigh et al., 2018). These studies demonstrate that single cell transcriptome profiling can offer a more nuanced view of the blastema not possible with genetic lineage analyses alone. Addressing similar questions in the context of mouse digit tip regeneration is important, especially in the context of working towards regenerative therapies.

In this paper we build upon previous findings that the mouse digit tip blastema is heterogeneous and lineage restricted by generating single cell transcriptomes of four stages of regenerating mouse digit tip blastemas as well as unamputated control digit tips. We sequenced over 38,000 total cells, allowing us to comprehensively define the cell type heterogeneity of the blastema throughout regeneration. We analyze an integrated data set from all regenerative and control time points and find that a clear signature of the cell types found in the quiescent control digit tip already exists in the early blastema, supporting lineage restriction. We find that blastema population dynamics vary by cell type and we focus specifically on a population of fibroblasts enriched in early blastema stages as compared to unamputated control digit tips. Differential expression analysis concentrated on these blastema-enriched fibroblasts reveals ten highly significant genes. Of these, *Mest* is expressed broadly in the blastema by RNA in situ hybridization, but not in the unamputated digit tip. This finding supports the notion of a regeneration-specific factor and opens the door to more subtle transdifferentiation relationships within the fibroblast lineage. Collectively, this data has important implications for regeneration of other musculoskeletal tissues and our broad understanding of epimorphic regeneration in mammals.

## RESULTS

### The early blastema is heterogeneous in cell type

Previous studies have used tissue/cell-type specific mouse genetic lineage analyses to characterize the regenerating digit tip blastema as both cellularly heterogeneous and lineage restricted (Lehoczky et al., 2011; Rinkevich et al., 2011). However, these studies leave room for additional insights into the origin of the blastema cells, the complexity of the cellular heterogeneity, and lineage relationships within germ layers. Toward these questions, we set out to characterize the adult mouse digit tip blastema as it first emerges from the stump tissue. By histology in outbred mice, the blastema is first detected at 10dpa (Fernando et al., 2012). While this timing is consistent with inbred FVB/NJ mice used in our study, we find gross microdissection of the blastema is not possible until 11dpa due to lack of tissue integrity at earlier stages. We amputated adult FVB/NJ mouse hindlimb digits 2, 3, and 4, midway through the terminal phalanx at a level permissive for innate regeneration (Figure 1A) (Fernando et al., 2012; Han et al., 2008; Neufeld and Zhao, 1995). At 11dpa we euthanized the mice and manually dissected the blastemas away from the surrounding epithelium and stump tissue. 12 blastemas dissected from two mice were pooled, dissociated, and subjected to single cell RNA sequencing using the 10X Genomics platform (Figure 1A). 7,830 cells were captured, with an average of 15,491 sequencing reads per cell. Quality control and filtering of reads was performed using a standard computational pipeline in Seurat, leaving RNA sequencing data for 7,610 high quality cells (Butler et al., 2018; Stuart et al., 2019), which was then used for unbiased cell clustering based on differential gene expression.

**Figure 1.**
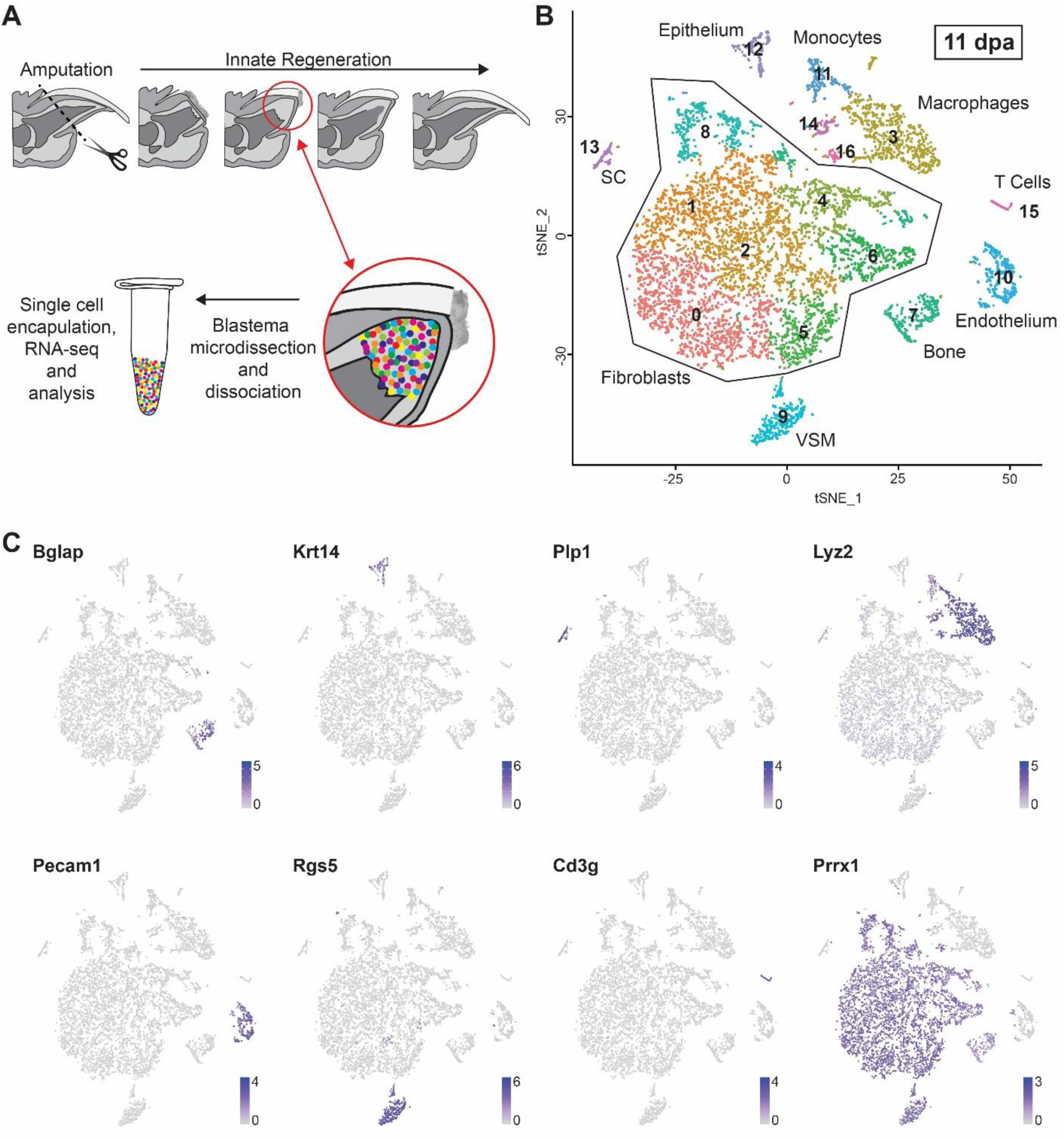
Cellular heterogeneity of the 11dpa blastema. (A) Schematic overview of innate mouse digit tip regeneration following amputation mid-way through the terminal phalanx. Schematic of the experimental design whereby blastemas were dissected, dissociated, and single cells captured. Single cell RNA libraries were prepared and sequenced for computational analysis. (B) Unbiased single cell clustering of 7,610 high quality cells visualized by tSNE plot. Each dot represents a single cell and cells assigned to the same cluster are similarly colored. Cell type identities are assigned as follows: fibroblasts (clusters 0-2, 4-6, and 8), macrophages (clusters 3, 14, and 16), bone (cluster 7), vascular smooth muscle cells (VSM) (cluster 9), endothelial cells (cluster 10), monocytes (cluster 11), epithelial cells (cluster 12), Schwann cells (SC) (cluster 13), T cells (cluster 15). (C) Gene expression tSNE overlay with examples of highly expressed, cell type specific markers used to assign cluster cell identities: *Bglap* (bone), *Krt14* (epithelial cells), *Plp1* (SCs), *Lyz2* (macrophages), *Pecam1* (endothelial cells), *Rgs5* (vascular smooth muscle cells), *Cd3g* (T cells), *Prrx1* (fibroblasts). Gray depicts low expression and purple depicts high expression as specified on the scale for each gene.

Unbiased clustering of the 11dpa blastema cells revealed 17 discrete populations (Figure 1B). We assigned cell identities to each of these clusters based on the top 10-20 most differentially expressed genes associated with each cluster and the known expression of these genes based on the literature, as well as checking expression of broadly established cell type marker genes in each cluster (Supplemental Table 1A). The assigned cell types include: fibroblasts (clusters 0-2, 4-6, and 8; *Prrx1*, *Msx1*, *Vim*); bone (cluster 7; *Bglap, Ibsp, Spp1*); monocytes (clusters 11; *Lyz2, Cd14, Cd86*) and macrophages (clusters 3, 14, 16; *Adgre1, C1qa, Lyz2*); T cells (cluster 15; *Cd3g, Icos, Trdc*); endothelial cells (cluster 10; *Pecam1, Cd93, Egfl7*); vascular smooth muscle cells (cluster 9; *Rgs5, Notch3, Myh11*), Schwann cells (cluster 13; *Plp1, Mbp,* and *Scn7a*); and epithelial cells (cluster 12; *Krt14, Krt42, Perp*) (Figure 1B and 1C). Our experiment was designed to capture the cells within the blastema, so we interpret the presence of epithelial cells within our sample as a technical artifact secondary to dissection (Figure 1B, cluster 12). While the nail and wound epithelia are both important cell types necessary for digit tip regeneration (Fernando et al., 2012; Lehoczky and Tabin, 2015; Mohammad et al., 1999; Takeo et al., 2013) they were not intentionally captured in this study and have been excluded from all of our analyses. We also interpret the presence of a mature bone population at 11dpa as a dissection artifact consistent with inclusion of stump bone adjacent to the blastema (Figure 1B, cluster 7). We include this cluster in our analyses as these cells provide terminally differentiated tissue to facilitate analysis of osteoprogenitor differentiation in the blastema.

Of the non-epithelial 11dpa populations, we captured several cell types already described in mouse digit tip regeneration. We find most cells in our sample are fibroblasts marked in particular by *Prrx1* and *Msx1*; previous genetic lineage analyses with these markers demonstrate that these cell types contribute broadly to the blastema (Lehoczky et al., 2011; Rinkevich et al., 2011). In addition, de-differentiated Schwann cells have been shown to secrete growth factors that may play a role in expansion of blastema cells during regeneration (Johnston et al., 2016), and in line with this we observe a population of Schwann cells (Figure 1B and 1C, cluster 13). Macrophages have been described in the post-amputation digit tip during wound closure and have been shown to be necessary for successful digit tip regeneration (Simkin et al., 2017). While this previous study finds peak numbers of macrophages prior to blastema formation by histology (Simkin et al., 2017), our blastema stage single cell analysis identifies three discrete macrophage populations (Figure 1B, clusters 3, 14, and 16), with one of the populations (cluster 14) likely representing mitotic macrophages based on expression of cell cycle genes such as *Top2a* and *Cdk1.* In addition, we find a population of endothelial cells (Figure 1B, cluster 10). *Sca1*/*Ly6a* positive endothelial cells have been characterized in the 10dpa blastema (Yu et al., 2014) and in line with previous data, 68% of the 11dpa endothelial cells in our dataset express *Sca1*/*Ly6a* (Supplemental Table 1A). Collectively, the presence of these previously described populations (Schwann cells, macrophages, endothelial cells, and fibroblasts) validates the robustness of our experimental approach. Importantly, unbiased single cell RNA sequencing also enabled us to identify cell populations that have not formally been described in the digit tip blastema. We isolated vascular smooth muscle cells and a small population of T cells (Figure 1B, clusters 9, and 15). We also isolated monocytes which likely contribute to the local macrophage population (Figure 1B, cluster 11).

To begin exploring the relationships among these populations, we constructed a cluster dendrogram and find the clusters fall into four main branches. The bone cells (cluster 7) make up their own branch of the dendrogram. Monocytes (cluster 11) and two macrophage populations (clusters 3 and 14) make up a separate branch (Supplemental Figure 1A), which correlates with the known lineage relationship between monocytes and macrophages (van Furth and Cohn, 1968; Virolainen, 1968). A third branch of the dendrogram is comprised of the fibroblast populations (clusters 0-2, 4-6, and 8) and the remaining macrophage population (cluster 16). While we expected the fibroblast populations to be closely related, the presence of a macrophage population in this clade was unanticipated. The fourth branch is made up of the remaining unrelated populations: T-cells, endothelial cells, vascular smooth muscle cells, epithelial cells, and Schwann cells. As validation of the cell population relationships, we calculated the Pearson correlation between in silico bulk transcriptomes of each cluster (Supplemental Figure 1B). Consistent with the dendrogram, the resultant correlation matrix shows that the fibroblast clusters are highly correlated with macrophage cluster 16. Many of the genes marking cluster 16 indicate that it is made up of macrophages (*C1qa* (83%)*, Adgre1* (74%)), while this cluster also expresses fibroblast marker genes (*Prrx1* (100%)*, Fmod* (100%)). While this could be a rare hybrid cell type, the mixed expression is more parsimonious with doublet cells (encapsulating two cells in one droplet before library preparation). To investigate this, we analyzed our dataset with DoubletFinder (McGinnis et al., 2018) and found 419 cells (5.5 %) classified as potential doublets, including 41 out of 53 cells (77%) in macrophage cluster 16 (Supplemental Figure 2, 11dpa). This finding prompted us to remove all putative doublet cells from subsequent analyses, though the existence of hybrid cell types has not been formally ruled out.

Although fibroblasts are known to participate in digit tip regeneration (Lehoczky et al., 2011; Marrero et al., 2017; Rinkevich et al., 2011; Y. Wu et al., 2013), the high proportion of the 11dpa blastema comprised of fibroblasts and the heterogeneity of these cells is striking, and has not been described previously. To begin to understand the biological significance of the seven discrete 11dpa fibroblast populations, we investigated the cluster marker genes that specifically mark these populations (Supplemental Table 1A). While all fibroblast clusters have gene expression in common, such as *Prrx1, Msx1,* and *Pdgfra* (Carr et al., 2018; Lehoczky et al., 2011; Rinkevich et al., 2011), these broad fibroblast markers ultimately mask the underlying heterogeneity of fibroblasts in the blastema. In line with the dendrogram and correlation matrix, clusters 0, 1, and 2 show common expression of many genes like *Ndnf*, *Matn4*, and *Mest* (Supplemental Figure 1C). However, *Ccl2* expression is more specific to cluster 0, and *Mmp13* expression to cluster 1, both perhaps consistent with a role in cytokine signaling or immune response (Supplemental Table 2). Cells in clusters 4 and 6 are not closely related and have distinct expression profiles, for example *Acan* and *Scara5* respectively, and GO analysis predicts involvement in the different biological processes of skeletal development and ECM organization for cluster 4 and iron ion import and transmembrane receptor protein tyrosine kinase signaling pathway for cluster 6 (Supplemental Figure 1C and Supplemental Table 2). Cluster 5 is also predicted to be involved in ECM organization, though these cells also express different genes than cluster 4, including *Aldh1a2*. Cluster 8, which expressed markers of proliferation such as *Top2a,* is comprised of mitotic fibroblasts. Taken together this demonstrates the heterogeneity of fibroblastic cells in the blastema, which suggests that blastema fibroblasts may participate in a diverse set of functions and lineages in the regenerating digit tip.

### Signature of the terminally differentiated digit tip already exists in the early blastema

The diversity of cell types we find by single cell RNAseq in the 11dpa blastema is supported by previous studies which demonstrate the mouse digit tip blastema is heterogeneous through genetic lineage analyses and staining for cell type specific markers (Carr et al., 2018; Johnston et al., 2016; Lehoczky et al., 2011; Lehoczky and Tabin, 2015; Rinkevich et al., 2011). However, digit tip regeneration is a prolonged, dynamic process, and little is known about how the heterogeneous blastema resolves into regenerated tissues or how the blastema cells relate to the cells of the original digit tip. Toward addressing these questions, we generated single cell RNAseq data from progressive blastema stages, as well as from the mesenchyme of unamputated digit tips. As with our 11dpa experiment, we amputated adult FVB/NJ mouse hindlimb digits 2, 3, and 4 (Figure 1A) and manually dissected blastemas at 12, 14, or 17dpa. For unamputated samples, mice were euthanized and non-epithelial tissues distal to our standardized amputation plane were dissected from hindlimb digits 2, 3, and 4. All four samples were separately dissociated and subjected to single cell RNA sequencing as above. Likely due to variation in cell dissociation, encapsulation, and library preparation we captured a range of cell numbers and reads for our samples: 12dpa (3,433 cells/27,628 average reads per cell), 14dpa (6,065 cells/27,026 reads), 17dpa (9,112 cells/21,416 reads), unamputated (UA) (13,750 cells/9,831 reads). Samples and reads were processed as with the 11dpa sample, and quality control and filtering left 3,309, 5,896, 8,778, and 12,871 cells in each data set respectively. We first analyzed each sample separately to determine which cell types were present at each regenerative stage (Supplemental Figures 3-6; Supplemental Tables 1B-1E). Intriguingly, all the cell types identified in the 11dpa blastema are also present in all four more mature blastema stages, as well as in the unamputated digit tip. Moreover, there are only a few additional cell types that appear in any sample and are limited to pre-osteoclasts (12dpa and 14dpa), neutrophils (14dpa), and a second population of Schwann cells (UA), though the emergence of these cells types in only certain regenerative stages could be explained by the differing number of sequenced cells.

The finding that the majority of cell types identified in the unamputated digit tip are already present in the 11dpa blastema presents at least two scenarios: 1) these are the same cells in terms of gene expression and only differ in quantity and perhaps spatial organization, or 2) these are cells within the same tissue-specific lineage that differ in gene expression and differentiation state at the time points sampled. Importantly, these possibilities need not be mutually exclusive given many of our assigned cell types have multiple discrete populations of cells (for example macrophages or fibroblasts) which could have separate roles in digit tip regeneration. To begin to address these questions, we removed all predicted doublet cells from the 11dpa, 12dpa, 14dpa, 17dpa, and UA single cell RNAseq data sets (Supplemental Figure 2), and combined and normalized the data for the remaining cells using the Integration workflow in Seurat (Stuart et al., 2019). Unbiased clustering of this combined data set revealed that cells from all stages were qualitatively well-mixed among 23 clusters (Figure 2A and 2B, Supplemental Table 3). This combined data set also allowed for increased resolution of cell types that might be rare in each stage and we now observed defined populations for myelinating (cluster 20, marked by *Mbp* and *Plp1*) and non-myelinating Schwann cells (cluster 14, marked by *C4b* and *Scn7a*), lymphatic endothelium (cluster 19, marked by *Pdpn* and *Lyve1*), and mitotic vascular smooth muscle cells (cluster 22, marked by *Rgs5* and *Top2a*) (Figure 2B). Overlaying each regenerative stage individually over the total integrated data set reveals that no cluster is comprised of cells from a single time point (Figure 2C), reinforcing the conclusion that unamputated digit cell types exist at all blastema stages and ruling out a broadly multipotent cell in the blastema.

**Figure 2.**
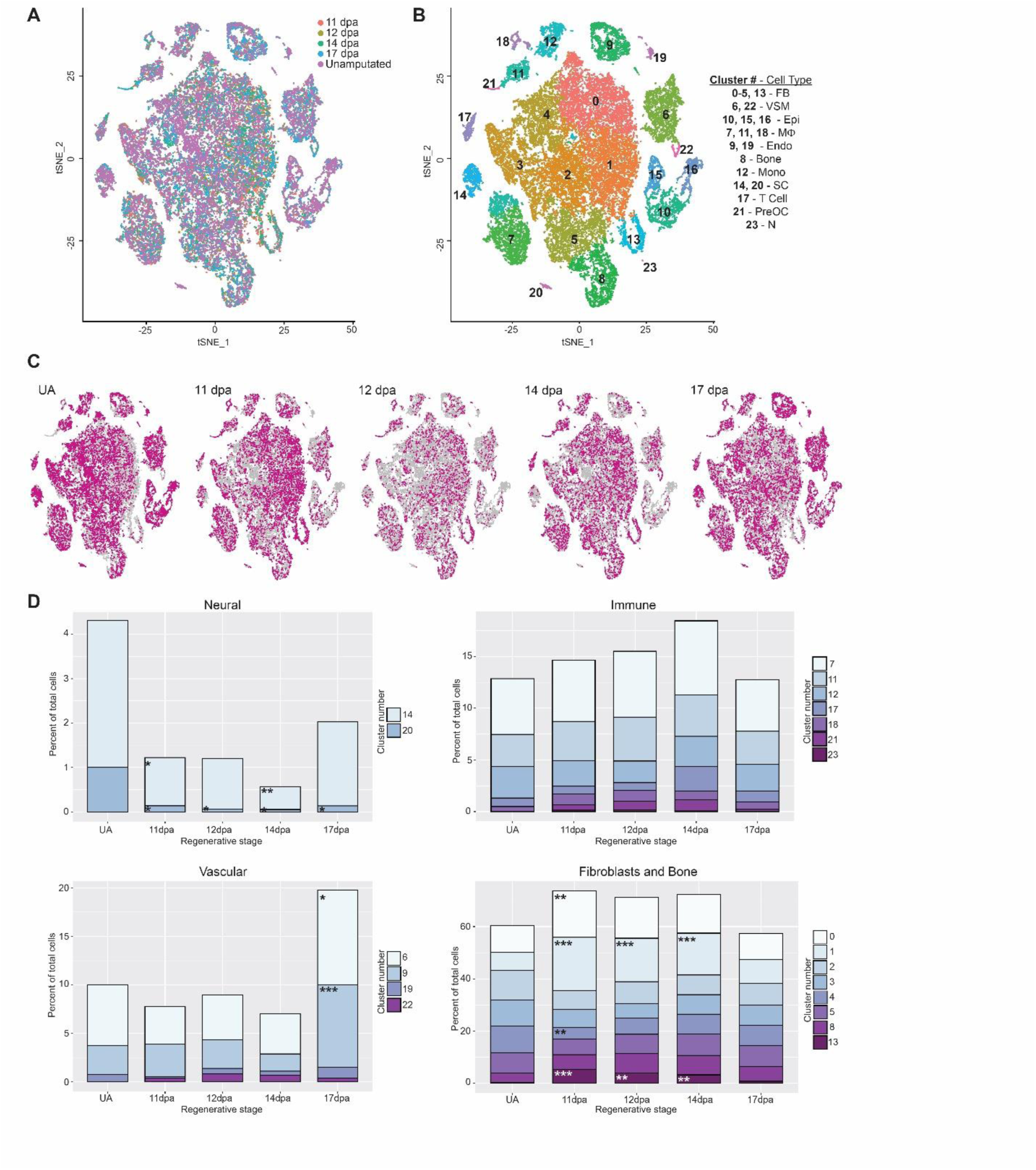
Integrated analysis of single cell populations through a regenerative time course. All analyses use combined and normalized 11dpa, 12dpa, 14dpa, 17dpa, and unamputated (UA) scRNAseq data sets. (A) tSNE plot of integrated data sets colored by regenerative stage: 11dpa (orange), 12dpa (olive green), 14dpa (green), 17dpa (blue), and unamputated (purple). (B) tSNE of integrated data sets showing clusters and cluster cell type annotations. Assigned cell types are: fibroblasts (FB; clusters 0-5, and 13), vascular smooth muscle cells (VSM; clusters 6 and 22), epithelial cells (Epi; clusters 10, 15, and 16), macrophages (Mф; clusters 7, 11, and 18), endothelial cells (Endo; clusters 9 and 19), bone (cluster 8), monocytes (Mono; cluster 12), Schwann cells (SC; clusters 14 and 20), T cells (cluster 17), pre-osteoclasts (PreOC; cluster 21), and neutrophils (N; cluster 23). (C) tSNE plot of integrated data set (gray) showing the cluster distribution of cells from each regenerative stage (pink). (D) The percentage of total cells represented by each cluster for the given regenerative stage. Each stage has been compared to the proportion of cells in UA, and significant changes were determined by differential proportion analysis (marked with asterisk). Clusters are categorized by overarching cell types (fibroblast or bone, immune, vasculature, or neural). Significance values are as follows: * denotes p<0.05, ** denotes p<0.01, *** denotes p<0.001.

### Blastema population dynamics during regeneration vary by cell type

To examine the dynamics of individual blastema cell populations throughout regeneration, we sought to assess 1) the proportion of cells in each cluster present at each regenerative stage and whether it changes over time, and 2) cell type specific gene expression changes through regeneration and whether this reflects tissue specific differentiation states. To compare cluster membership over time, we performed differential proportion analysis (Farbehi et al., 2019) on our integrated data set segregated by stage. This analysis allows for building hypotheses about the timing and function of blastema populations and whether they are regeneration or homeostasis specific. As a first pass analysis we looked for changes in relative population size as compared to the unamputated digit and found significant regenerative population dynamics for Schwann cells (clusters 14 and 20), vascular smooth muscle cells (clusters 6 and 9), and several fibroblast populations (clusters 0, 1, 4, and 13). Notably, no significant population dynamics were found in the immune-related clusters (Figure 2D, Supplemental Table 4).

Little is known about the influence of immune cells in digit tip regeneration. Only macrophages have been characterized and found to be necessary (Simkin et al., 2017), but it is important to understand whether additional immune cells play a role in the blastema as well as the initial inflammation response. In our data, differential proportion analysis finds no significant differences in proportion of monocytes, macrophages, pre-osteoclasts, T cells, or neutrophils between any two stages in our data set (Figure 2D, Supplemental Table 4). Given the small number of cells (565 total) in the pre-osteoclast, T cell, and neutrophil clusters, we are likely statistically underpowered to make meaningful conclusions for these cell types. That said, we have relatively large populations of macrophages and monocytes at all stages of our data (4,611 cells total), thus the absence of significant population dynamics for these cell types is likely reflective of the biology of the digit tip regeneration immune response. To understand the lineage relationship of these cells, and if there is a differentiation trend during regeneration, we subjected the cells in these clusters to SPRING force-directed trajectory analysis (Weinreb et al., 2018). The data reveals a major differentiation trajectory from monocytes to macrophages, with no skewing in differentiation state based on regenerative/unamputated stage (Supplemental Figure 7A-C). This finding suggests that the production of macrophages from monocytes in the digit tip is at a homeostatic rate once the blastema is formed. No specific lineage relationships are revealed for the population of ECM producing macrophages, T-cells, or neutrophils, though we have a minimal sampling of these populations (Supplemental Figure 7A). However, there is a qualitative increase in differentiation of monocytes to pre-osteoclasts and an increase in proliferative macrophages marked by *Adgre1* and *Top2a* in the blastema (Supplemental Figure 7B, 7D, and 7E). The presence of proliferative macrophages could reflect a lingering response to the initial wound or a physiological role in the blastema itself that is not satisfied by recruited monocytes.

Previous studies demonstrate that innervation and neural associated cell types such as Schwann cell precursors are necessary for digit tip regeneration (Carr et al., 2018; Dolan et al., 2019; Johnston et al., 2016; Mohammad and Neufeld, 2000; Takeo et al., 2013). Both sensory and sympathetic axons innervate the connective tissue of the unamputated digit tip, and they are accompanied by both myelinating and non-myelinating Schwann cells (Dolan et al., 2019). In the unamputated digit tip, we find that myelinating and non-myelinating Schwann cells make up 0.40% and 1.9% of the captured cells, respectively. Despite small numbers of cells, differential proportion analysis reveals a significant depletion of both populations in the 11dpa blastema compared to the unamputated digit tip (Figure 2D, Supplemental Table 4). Notably, the myelinating Schwann cells remain significantly reduced through all of our assayed stages and do not reinstate pre-amputation levels by 17dpa. This observation is in line with previous work, which showed that Schwann cells are present in the blastema but are qualitatively less abundant than in the quiescent digit tip, and only non-myelinating Schwann cells appear to recover to pre-amputation levels by 4 weeks post amputation (Dolan et al., 2019; Johnston et al., 2016). The reduction in population size of non-myelinating Schwann cells persists through 14dpa and begins to recover to pre-amputation levels by 17dpa. While it is important to understand whether these dynamics correlate with the differentiation trajectory of these cell types, these clusters (clusters 14 and 20) contained too few cells for a meaningful SPRING trajectory analysis, thus these questions remain for future experiments designed to specifically enrich these populations.

In previous studies, *VE-cadherin*-expressing endothelial cells have been shown to be lineage restricted during digit tip regeneration (Rinkevich et al., 2011) and individual endothelial cells are found in the blastema (Fernando et al., 2012). However, the overall dynamics of vascular-related cells in the blastema, including vascular smooth muscle cells, has not yet been characterized. Differential proportion analysis of our all stage integrated single cell RNAseq data reveals no significant change in the relative population sizes of endothelial cells or vascular smooth muscle cells between 11dpa or 14dpa compared to UA. However, at 17dpa, both populations are significantly expanded (Figure 2D). The low relative percentage of vascular cell types in the early regenerative stages is consistent with previous work that describes minimal angiogenesis in the early blastema (Yu et al., 2014), and the spike in vascular cell types at 17dpa could be indicative of over-sprouting of blood vessels before they are pruned (reviewed in Korn and Augustin, 2015). To explore these cell types further, we determined the differentiation trajectory using SPRING. The four vascular-related cell clusters (Figure 2B clusters 6, 9, 19, and 22) appear separate in the SPRING visualization, with a lineage of proliferative vascular smooth muscle cells streaming into the main cluster of vascular smooth muscle (VSM) cells. No major lineage relationship is found between vascular and lymphatic endothelial cells (Figure 3A). Lymphatic endothelial cells have not yet been described in digit tip regeneration so it is an important advance to have captured them, and their associated markers; that said, this is an extremely small population and we are underpowered to make conclusions on a differentiation trajectory. When analyzing the VSM cells and the vascular endothelial populations by regeneration stage, there is qualitative spatial variation between the different time points on the SPRING plot (Figure 3B). Closer evaluation of the VSM cells confirms that all of the cells express the tissue-specific marker *Rgs5* (Figure 3C) (Li et al., 2004). However, early blastema cells (11, 12, and 14dpa) are concentrated on one side of the cluster, while UA cells are on the other and 17dpa cells appear throughout. This observation is consistent with the differentiation of VSM cells from a progenitor state to terminally differentiated cells and is supported by the differential expression of *Gadd45b* and *Lgals1* (Figure 3B and 3C) (Gizard et al., 2008; Kim et al., 2014; Moiseeva et al., 2000). Similarly, the cluster of vascular endothelial cells all express the broad marker *Pecam1* (Figure 3D) (Albelda et al., 1990; Müller et al., 2002), though the trajectory shows UA cells concentrated on one edge and blastema cells throughout the rest of the cluster, consistent with differentiation from vascular endothelial progenitors (expression of *Egfl7* (Campagnolo et al., 2005)) to terminally differentiated cells (expression of *Rnd1* (Suehiro et al., 2014)) (Figure 3D). Overall, the different vascular populations have recovered by 17dpa, and there are differences in differentiation state between the blastema stages and the unamputated digit tip. The 17dpa vascular cells span both mature and de-differentiated states.

**Figure 3.**
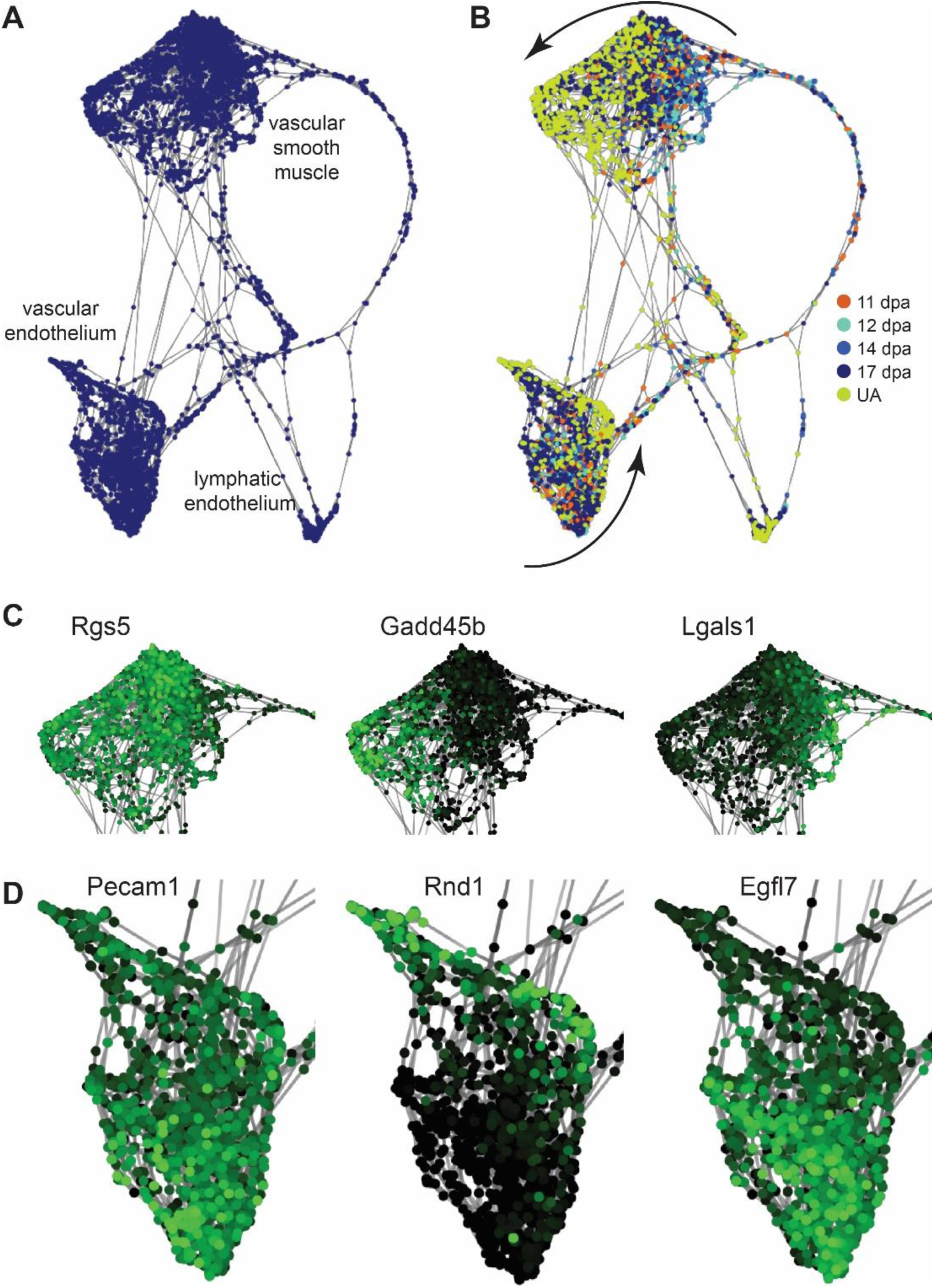
Vasculature differentiation trajectory of integrated data set. SPRING lineage trajectory analysis of cells from the integrated data set vascular clusters 6, 9, 19 and 22. (A) Force-directed plot of cells showing clusters of vascular smooth muscle cells, vascular endothelial cells, and lymphatic endothelial cells. (B) SPRING plot as in (A) with regenerative stages of each cell colored coded: 11dpa (orange), 12dpa (light blue), 14dpa (medium blue), 17dpa (dark blue), unamputated (yellow). Differential clustering of blastema cells and unamputated cells suggests tissue specific differentiation of the vascular smooth muscle cells and the vascular endothelium (curved arrows). (C) Gene expression overlay on vascular smooth muscle cells. *Rgs5* is expressed in all cells, *Gadd45b* is more highly expressed in UA cells, and *Lgals1* is more highly expressed in blastema cells. High expression is in green and low expression is black. (D) Gene expression overlay on vascular endothelial cells. *Pecam1* is expressed in all cells, *Rnd1* is more highly expressed in UA cells, and *Egfl7* is more highly expressed in blastema cells.

### Diversity and dynamics of fibroblasts during regeneration reveal regeneration-specific markers

As fibroblasts make up the majority of the blastema and are more heterogeneous than previously described (Supplemental Figure 1) (Lehoczky et al., 2011; Rinkevich et al., 2011; Y. Wu et al., 2013), we analyzed all fibroblast and bone cells separately from the rest of the cell types. This unbiased clustering resulted in 15 populations that were broadly concordant with the original all-cell-type clustering, yet more refined (Figure 4a compared to Figure 2a). We performed SPRING analysis on the all-cell-integrated data set and found no populations transdifferentiating from fibroblasts or bone into any other cell type within the blastema or unamputated digit tip (Figure 4B), supporting the lineage restriction found in previous genetic lineage studies (Lehoczky et al., 2011; Rinkevich et al., 2011). Among the fibroblast clusters there is a distinct differentiation trajectory from clusters 1 and 7 into bone (cluster 8; *Bglap* and *Ibsp* expression) (Figure 4C). While the presence of mature bone cells at all regenerative stages (Figure 4G) is an artifact of our microdissection, these cells facilitate trajectory mapping and allow for cell type identification of clusters 1 and 7 as osteoprogenitors and differentiating osteoblasts, respectively (Figure 4C, *Postn* expression; Supplemental Table 5). Another possible group of trajectories originate in cluster 2, then branch and terminate in clusters 5, 6, 9, 12, and 14 (Figure 4D). It is unclear if these trajectories reflect the differentiation of resident fibroblast subtypes within the digit tip, or whether they reflect skeletal tissue lineages (ex. tenocytes or adipocytes). Clusters 6 and 12 express several tendon-specific genes, such as *Fibin* and *Tnmd* (Supplemental Table 5) (Brandau et al., 2001; Pearse et al., 2009), and we hypothesize that cluster 2 contains mesenchymal stem cells (MSCs) or minimally, tendon progenitor cells because of *Scx* expression (Schweitzer et al., 2001). However, no discrete skeletal lineage can be assigned to clusters 5, 9, and 14 based on marker gene expression, for example *S100a4* and *Smoc2*, thus they may be incompletely differentiated MSCs that also reside in the unamputated digit, or resident fibroblast subtypes that have not been characterized. Clusters 0, 3, and 4 make up a third major concentration of cells. They do not appear to differentiate into a specific lineage and remain centrally located on the trajectory map (Figure 4F). Intriguingly, this analysis reveals that these clusters are enriched for cells from early blastema stages and this is not a function of proliferation (Figures 4E and 4G). This finding could be consistent with the dedifferentiation of fibroblasts (lineage contribution), or regenerative-specific fibroblasts (non-lineage, providing signals).

**Figure 4.**
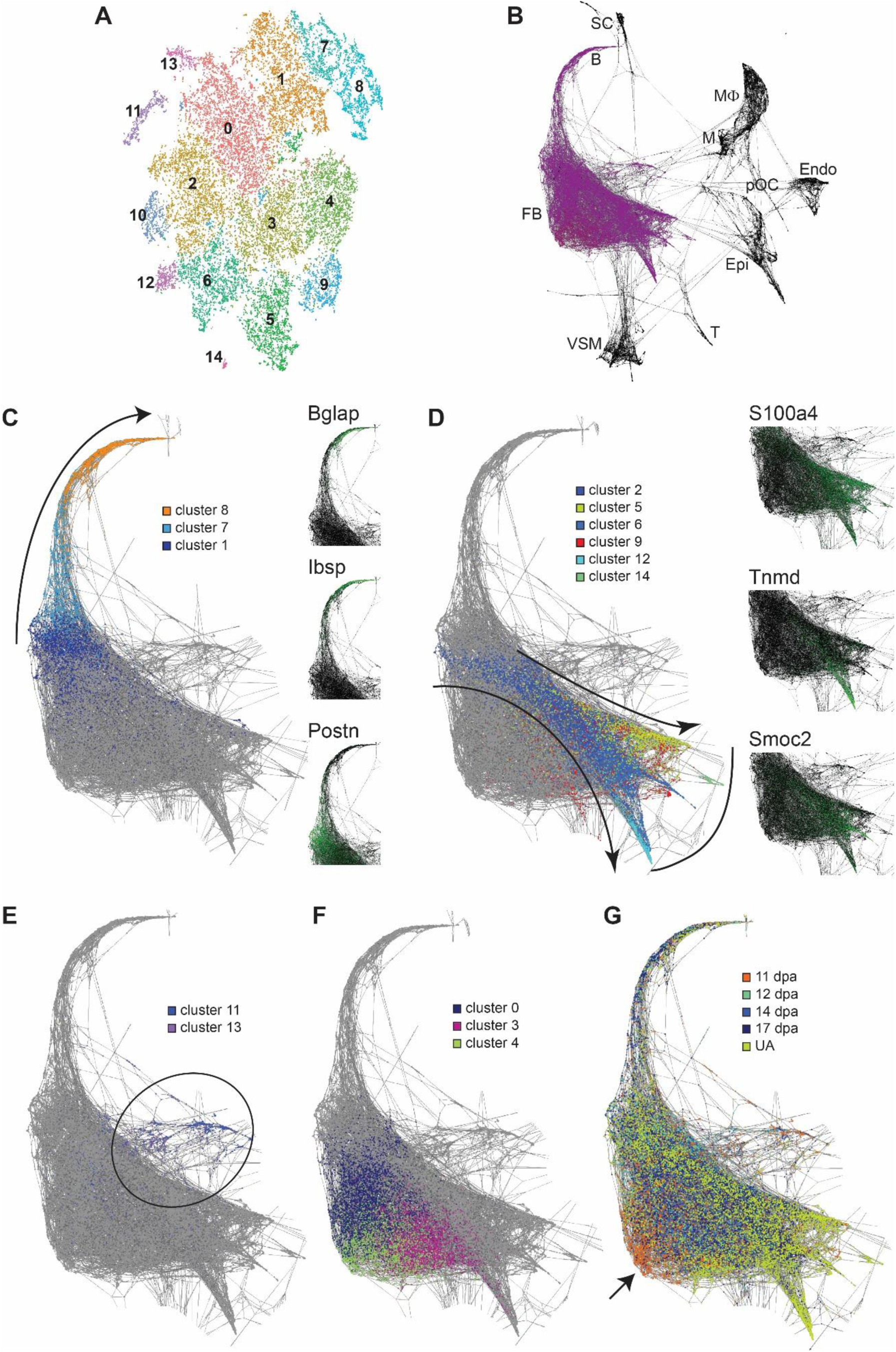
Fibroblast differentiation trajectory of integrated data set. (A) tSNE plot of unbiased re-clustering of fibroblast and bone cells from integrated data set (Figure 2: clusters 0-5, 8, and 13), reveals 15 refined clusters. (B) SPRING lineage trajectory analysis of cells from the integrated data set showing fibroblasts (FB) or bone (B) do not transdifferentiate into Schwann cells (SC), monocytes (M), macrophages (MΦ), pre-osteoclasts (pOC), endothelium (Endo), epithelium (Epi), T cells (T), or vascular smooth muscle (VSM). Fibroblast SPRING lineage trajectory overlaid with (C) bone lineage from cluster 1 to cluster 8 (curved arrow). Marker gene expression for each cluster shown with *Bglap*, *Ibsp*, and *Postn*. High expression is in green and low expression is black. (D) Proposed mesenchymal stem cell lineage from cluster 2 to clusters 5, 6, 9, 12, and 14 (curved arrows), with distinct lineages marked by *Tnmd*, *S100a4*, and *Smoc2*. Curved line depicts terminally differentiated cells. (E) Clusters 11 and 13 mark mitotic cells (black circle) and (F) clusters 0, 3, and 4 do not contribute to a lineage, but are (G) enriched for early stage blastema cells (arrow pointing to orange).

Differential proportion analyses of the re-clustered fibroblasts support our qualitative findings from the trajectory analysis. Clusters 1, 3, 7, 8, 9, and 14 do not have significant changes in population size through regeneration, though for clusters 1, 7, and 8, this can be attributed to inclusion of bone in all dissections (Figure 5A and Supplemental Table 6). Cell populations in clusters 2, 5, 6, and 12 all are significantly depleted at 11dpa, and are restored to unamputated levels by 17dpa, with the exception of cluster 12 (Figure 5B and Supplemental Table 6). This profile may be consistent with amputated tissue lineages being restored through regeneration. Unexpectedly, we found the cell populations in clusters 0, 4, 10, 11, and 13 to be significantly increased at 11dpa; by 17dpa clusters 0 and 10 have still not returned to unamputated levels (Figure 5C and Supplemental Table 6). For cluster 11 and 13, these population dynamics can be attributed to cellular proliferation (Figure 4E and Supplemental Table 5), however for clusters 0, 4, and 10 this suggests a regeneration specific function. Gene expression analysis between cells from blastema-enriched clusters (Figure 5C) and blastema-depleted clusters (Figure 5B) results in 370 significantly differentially expressed genes (Supplemental Table 7). We prioritized these genes by selecting those with an average log fold-change ≥ 0.75 and with the percent of cells in other clusters expressing the gene ≤ 0.25, leaving 10 genes (Figure 5D). Of these, several had distinct regeneration-specific expression profiles. *Ccl2* and *Cxcl2* both showed increased expression at early blastema stages, with low expression in late regeneration as well as the unamputated digit tip (Figure 5E). *Mmp13* and *Mest* both showed expression at all blastema stages, with negligible expression in the unamputated digit tip (Figure 5E). While *Mmp13* has already been implicated in regeneration in other species as a necessary mediator of ECM remodeling (Calve et al., 2010; Miyazaki et al., 1996; Vinarsky et al., 2005; C.-H. Wu et al., 2013), *Mest* is a novel marker of the blastema and epimorphic regeneration.

**Figure 5.**
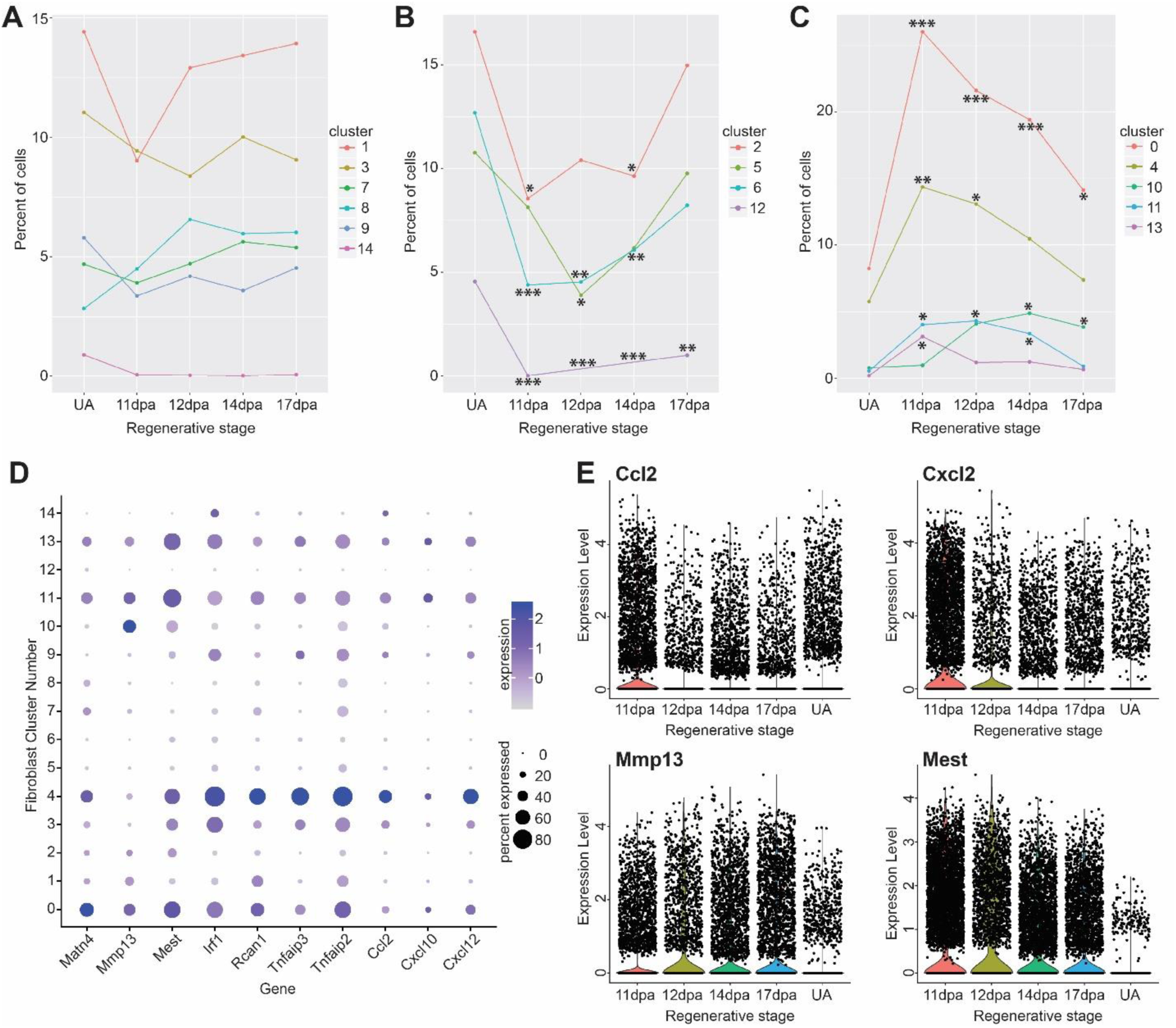
Analysis of blastema fibroblast population dynamics. Differential proportion analysis of fibroblast clusters parsed by regeneration profile where clusters in (A) have no significant population dynamics between blastema stages and unamputated. (B) Cells in clusters 2, 5, 6, and 12 are significantly depleted as compared to unamputated and (C) cells in clusters 0, 4, 10, 11, and 13 are enriched during regeneration as compared to unamputated. Significance values are as follows: * denotes p<0.05, ** denotes p<0.01, *** denotes p<0.001. (D) Subset of genes enriched in blastema stages as compared to unamputated. Gray depicts low expression and dark purple is high expression; small circles depict a low percentage of cells and large circles depict a high percentage. (E) Violin plots of representative genes enriched in blastema fibroblasts as compared to unamputated. Black points represent individual cells and the colored curve shows the distribution of cells at a given expression level.

To determine the distribution of *Mest* expressing cells within the regenerating mouse digit tip, we utilized RNA in situ hybridization. The *Mest* antisense RNA probe revealed the expected expression domains, including tongue and vertebrae, on control E12.5 embryonic mouse sections (Supplemental Figure 8A-C). No significant *Mest* expression was found on unamputated digit tip sections (Figure 6A) and appeared comparable to *Mest* sense RNA control probe hybridization on unamputated digits (Supplemental Figure 8E). In contrast, at 11dpa *Mest* expression is found scattered throughout blastema cells which is not seen for sense RNA probe 11dpa controls (Figure 6B and Supplemental Figure 8D). Heterogeneous *Mest* blastema expression becomes even more pronounced at 12dpa and 14 dpa, then begins to decrease and become centrally restricted at 17dpa (Figure 6C-E). These in situs validate our computational analysis and establish *Mest* as a novel regeneration-specific marker of mouse digit tip blastema.

**Figure 6.**
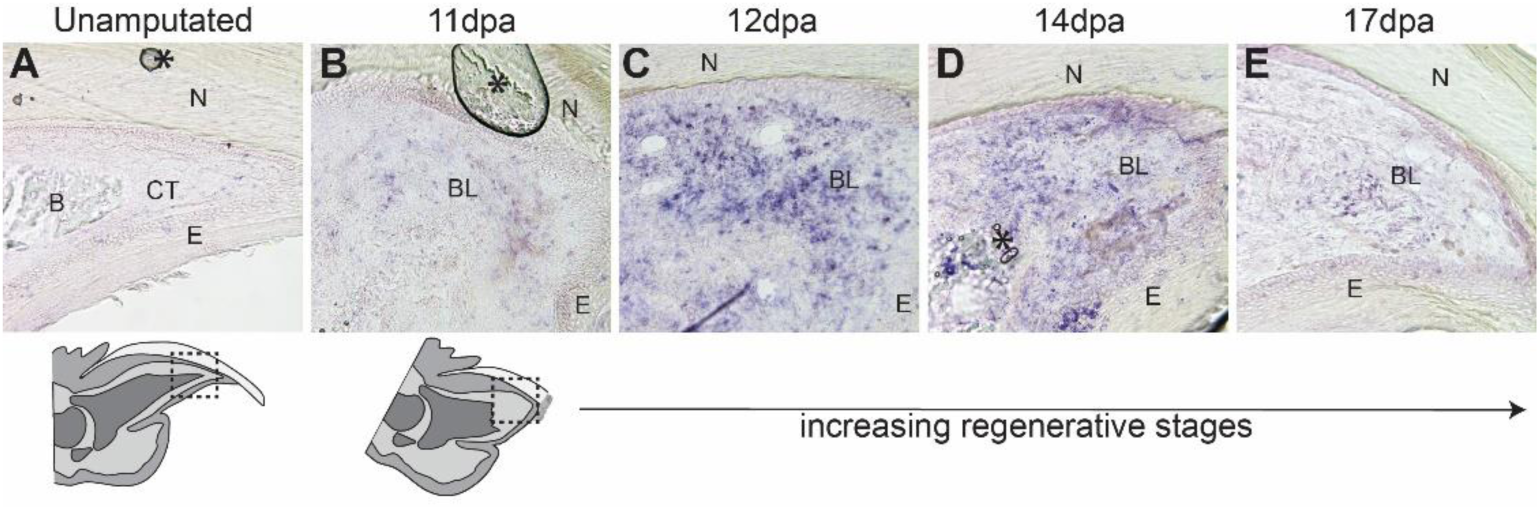
*Mest* expression during digit tip regeneration. RNA section in situ hybridization for *Mest* on regenerating digit tips. (A) Unamputated digit tip with orientation shown by schematic below. (B) 11dpa with region of the blastema depicted in the schematic below. Additional regenerative stages include (C) 12dpa, (D) 14dpa, and (E) 17dpa. Asterisks (*) denote artifacts from coverslipping. Abbreviations: (N) nail, (CT) connective tissue, (E) epithelium, (B) bone, (BL) blastema.

## DISCUSSION

Historically, the blastema has been described as a collection of proliferative and homogeneous cells that give rise to the regenerated tissue (Hay and Fischman, 1961). Based on this description, we would expect there to be a high proportion of actively dividing cells in the blastema. We find a dividing fibroblast cluster in 11, 12, and 14dpa blastema stages that is depleted by 17dpa and not found at all in the unamputated digit tip. These dividing fibroblasts account for less than 10% of the total cells captured, which is consistent with previous results using EdU (Johnston et al., 2016). This challenges the idea of the blastema as a highly proliferative structure and suggests that relatively few proliferative cells are needed to support regeneration once the blastema is formed.

The classical assessment of the blastema as homogeneous was based on cell morphology in the regenerating newt limb (Hay and Fischman, 1961), however recent studies of digit tip regeneration use genetic lineage tracing to collectively conclude that the digit tip blastema contains progenitors that are heterogeneous in cell-type and lineage restricted (Johnston et al., 2016; Lehoczky et al., 2011; Rinkevich et al., 2011). While these studies clearly demonstrate the blastema is not pluripotent across tissue germ layers, multipotency within germ layers was never formally addressed likely due to the tissue-specificity limitations of the available genetic alleles. Our single cell RNAseq analysis reveals that all of the defined cell populations of the unamputated digit tip are already present in the 11dpa blastema, validating both the lineage restriction and heterogeneity of the blastema as defined by genetic lineage analyses (Lehoczky et al., 2011; Rinkevich et al., 2011). This regenerative dataset allows for an unbiased view of the cell types within the mouse digit tip blastema, and includes several cell types that have already been described during digit tip regeneration including Schwann cells, macrophages, neutrophils, endothelial cells, osteoblasts, and fibroblasts (Johnston et al., 2016; Lehoczky et al., 2011; Rinkevich et al., 2011; Simkin et al., 2017). Importantly, our clustered data reveals sub-populations and more detailed gene expression associated with these previously reported populations, including two Schwann cell populations, three macrophage populations, and 15 distinct fibroblast populations (Figures 2B and 4A). Our data also provides insight into cell types that have not previously been described during digit tip regeneration, including T cells, monocytes, pre-osteoclasts, vascular smooth muscle cells, and lymphatic endothelium. From the standpoint of epimorphic regeneration, this adds considerable information to our understanding of the number of unique cell types participating in digit tip regeneration. From an experimental standpoint, we now have access to specific genetic markers for each of these individual cell types to refine future in vivo experimentation.

Integration of our longitudinal regenerative data set reveals that a signature of unamputated digit tip cell types exists in the early blastema. Importantly, these are not necessarily identical cell populations and can instead be related cells in distinct cell states (Morris, 2019). With trajectory analysis, we find differentiation from monocytes to macrophages equally at all regenerative stages, however we do find more blastema cells in the pre-osteoclast lineage than are found in the homeostatic digit tip (Supplemental Figure 7E). This likely indicates that our analysis missed the post-amputation macrophage response which occurs prior to the emergence of the blastema (Simkin et al., 2017). Conversely, our data finds discrete clusters of vascular related cells (vascular smooth muscle, vascular endothelium, and lymphatic endothelium) taking on distinct cell states throughout digit tip regeneration (Figure 3). Our analysis provides a refined view of these tissue-specific differentiating cells; for example, a canonical cell-type specific marker such as *Pecam1* would label all vascular endothelial cells, whereby our data details genes and timing of emergence of different populations potentially useful for experimental access to vascular endothelial progenitors (Figure 3, *Egfl7*) or terminally differentiated cells (Figure 3, *Rnd1*).

A similar analysis with the digit tip fibroblast and bone populations enriches our previous understanding of heterogeneity and lineage restriction within the connective tissue and skeletal lineages of the regenerating digit tip. The extensive fibroblastic heterogeneity seemed unprecedented given the limited number of mesenchymally-derived tissues within the digit tip regenerate, which includes bone and tendon but not cartilage or muscle. This may suggest that only a portion of the fibroblast populations are progenitors (mesenchymal stem cells (MSCs)) differentiating into tissue-specific lineages, whereby the remaining populations might function as niche fibroblasts for ECM production, chemotaxis, etc. Trajectory analysis with these cells indeed reveals multiple tissue-specific lineages, including osteoprogenitors into bone, as well as MSCs into tendon (Figure 4). From this analysis it is not clear if these progenitors can transdifferentiate between skeletal lineages, though it certainly seems possible (Figure 4, clusters 1 and 2). This analysis also underscores the importance of re-visiting conclusions from previous fibroblastic genetic lineage analyses, as it is likely that these cre alleles (ex. *Msx1* or *Prrx1*) mark the majority of our newly defined fibroblastic clusters, ultimately limiting the conclusions about transdifferentiation that can be drawn (Lehoczky et al., 2011; Rinkevich et al., 2011).

Beyond lineage restriction and heterogeneity, our data offers new insight into the molecular biology of digit tip regeneration. Differential gene expression analysis between blastema cells and homeostatic digit cells enabled us to identify markers of regenerating fibroblasts (Figure 5 and Supplemental Table 7). We found several such markers of blastemal fibroblasts that are upregulated in clusters associated with regeneration and not the quiescent digit tip, including some associated with inflammation (*Ccl2, Cxcl2*) and some that regulate extracellular matrix (*Matn4, Mmp13*). The gene with the most dramatic change in expression from unamputated digit tip to blastema is *Mest*. The molecular function of *Mest* is not known, but it bears resemblance to the α/β hydrolase family of enzymes and is important for embryonic growth (Lefebvre et al., 1998). Intriguingly, *Mest* has been associated with other regenerative models, in particular the regeneration of adipocytes and hair follicles following skin wounding, where it is thought to be a marker of de-differentiated fibroblasts that differentiate into myofibroblasts (Guerrero-Juarez et al., 2019). The role of *Mest* in digit tip regeneration needs to be explored in vivo. It will be important to determine whether *Mest*-expressing cells are MSCs or de-differentiated fibroblasts that can transdifferentiate into multiple mesenchymal lineages or whether these cells are regeneration-specific fibroblasts that do not contribute to a tissue lineage, but instead provide niche factors. These findings can give insight into inducing epimorphic regeneration in other mammalian tissues.

This work presents extensive new and refined data for the regenerating mouse digit tip. Moving forward, much experimental work is required to determine which of these cell types and genes are necessary for regeneration and what molecular role they play. Studies on the necessity and role of Schwann cells and macrophages exemplify the types of focused experiments needed to put this comprehensive digit tip cell atlas into biological context (Johnston et al., 2016; Simkin et al., 2017). Importantly, our study cannot conclusively define the origin of blastema cells and whether they arise via de-differentiation of terminally differentiated cells or whether they are derived from tissue-resident progenitor cells. Our data suggest that both could be true, depending on the lineage. For instance, macrophages in the blastema appear to originate from resident monocytes (Supplemental Figure 7B), whereby vascular cells and at least a subset of fibroblasts may de-differentiate to form the blastema (Figures 3B and 4G). Future experiments, taking advantage of the markers defined in this work, are needed to formally distinguish between these mechanisms for each cell-type.

## MATERIALS AND METHODS

### Mouse digit tip amputation surgery

All mice were housed in the Hale BTM specific pathogen free vivarium at Brigham and Women’s Hospital. All mouse breeding and surgery was performed in accordance with BWH IACUC approved protocols. All experiments used inbred wild-type FVB/NJ mice (JAX 001800), maintained in our colony. 6-week-old adult male mice were used for unamputated controls and digit tip amputation surgeries and subsequent blastema collection; 2 mice were used for each time point (12 total hindlimb digits). Mice were anesthetized with inhaled isoflurane (1-2% in oxygen) and digits were visualized with a Leica MZ6 stereomicroscope. For each mouse, digits 2, 3, and 4 of both hindlimbs were amputated midway through the distal digit segment using a #11 disposable scalpel. Subcutaneous buprenorphine (0.05 mg/kg) was given peri- and post-operatively as analgesia. Post-surgical animals were housed individually. Mice were euthanized and digits were collected at 11, 12, 14, and 17 days post amputation for blastema collection.

### Digit tip single cell isolation

For all regenerating digits, blastemas were microdissected from the digit tip while being visualized under a Leica M165FC stereomicroscope. To minimize collection of epithelial cells, the nail and associated epithelium was reflected and removed, leaving direct access to the blastema. The blastema was removed intact with super-fine forceps and placed into ice-cold PBS. Control unamputated digit tip samples were collected in a similar manner whereby the nail and associated epithelium was removed and the exposed digit tip bone and attached connective tissues were amputated with a #11 scalpel at a position comparable to all other digit tip amputations. These control digit tips were collected into ice-cold PBS and processed in parallel with the blastema samples. All tissues were enzymatically dissociated with trypsin (Thermo Fisher) (0.25%, 37°C for 1 hour), then with collagenase type I (Thermo Fisher) (0.65%, 37°C for 20 minutes), followed by manual trituration with a pipette. Red blood cells were lysed using ACK lysis buffer. Dissociated cells were washed, filtered, and resuspended in 0.4% BSA in PBS for cell counting on. All samples were adjusted to a concentration of 1,000 cells/uL for the single cell RNAseq pipeline.

### Single cell capture, library construction and next generation sequencing

All single cell RNAseq experiments used the 10x Chromium commercially available transcriptomics platform (10x Genomics Inc) implemented by the Brigham and Women’s Hospital Single Cell Genomics Core. Single cells were captured using the 10X system; the 12dpa blastema sample cDNA library was made with Single Cell 3’ v2 chemistry, and all other libraries (11dpa, 14dpa, 17dpa, and UA) were made with Single Cell 3’ v3 chemistry. Libraries were sequenced at the Dana Farber Cancer Institute Molecular Biology Core Facilities on the Illumina NextSeq 500 sequencing system.

### Single cell clustering and differential expression analysis

Computationally intensive portions of this research were conducted on the O2 High Performance Computing Cluster, supported by the Research Computing Group at Harvard Medical School (http://rc.hms.harvard.edu) using R version 3.5.1 (R Core Team, 2018). 10x Genomics Cell Ranger software (version 3.0.2) was used to convert raw BCL files to FASTQ files, align reads to the mouse mm10 transcriptome, filter low quality cells, and count barcodes and unique molecular identifiers (UMIs). The cell by gene matrices for each of the five datasets generated by Cell Ranger were individually imported to Seurat version 3.0 (Stuart et al., 2019), and cells with unusually high numbers of UMIs (possible doublets) or mitochondrial gene percent (possible dying cells) were filtered out (thresholding in Supplemental Table 8). Gene counts were normalized using the LogNormalize method and highly variable genes selected for downstream analysis (variable feature selection described in Stuart et al., 2019). Data was scaled and principal components selected and adjusted for each experimental group of cells for dimensional reduction (Supplemental Table 8). Cells were clustered using the standard Seurat workflow and visualized using t-distributed stochastic neighbor embedding (tSNE) (van der Maaten and Hinton, 2008). Cluster markers were found using FindAllMarkers with the Wilcoxon rank sum test, with only.pos = TRUE, min.pct = 0.25, logfc.threshold = 0.25. For the blastema-enriched vs. blastema-depleted differential expression analysis, FindMarkers was run on the fibroblast only Seurat object with clusters determined to be expanded in the blastema (0, 4, 10, 11, and 13) as ident.1 and clusters determined to be depleted in the blastema (2, 5, 6, and 12) as ident.2. All other parameters were default.

### Hierarchical and correlation analyses

The dendrogram of 11dpa cell populations was built in R version 3.5.1 (R Core Team, 2018) using the Seurat version 3.0 (Butler et al., 2018; Stuart et al., 2019) command BuildClusterTree on the 11dpa Seurat object with default parameters. The dendrogram was visualized using PlotClusterTree in Seurat. For the correlation analysis, bulk transcriptomes for each cluster were calculated using AverageExpression in Seurat. Pearson correlations were calculated from the resulting gene x cluster expression matrix using R base function cor with method = “pearson”. The correlation matrix was visualized using the corrplot function from the corrplot library (Wei and Simko, 2017).

### GO biological process category analysis

For GO category analysis of 11dpa fibroblast populations, cluster marker genes with adjusted p-value ≤ 0.05 and average log fold-change ≥ 0.5 were used as input to the PANTHER classification system web interface (http://pantherdb.org) (Mi et al., 2010; Thomas et al., 2003). The statistical overrepresentation test was used with the slim biological processes category, fisher’s exact test, and the Bonferroni correction for multiple hypothesis testing. All genes in the gene by cell matrix from the 11dpa Seurat object were used as the background set for the overrepresentation test.

### Cell doublet identification

Initial broad screening for doublets in each data set was performed via quality control processing in Seurat by UMI thresholding (Supplemental Table 8). For specific detection of putative doublet cells, we implemented the DoubletFinder (McGinnis et al., 2018) package in R version 3.5.1 (R Core Team, 2018) as described in detail at https://github.com/chris-mcginnis-ucsf/DoubletFinder. The doublet rate used was estimated from the 10x Chromium users guide and the number of cells captured, and is as follows: UA, 7.6%. 11dpa, 5.5%. 12dpa, 2.5%. 14dpa, 4.6%. 17dpa, 6.9%. All identified putative doublets were removed from data sets.

### Batch correction, dataset integration and sub-clustering

The cells for our five experimental samples were collected and processed on multiple days, potentially contributing to batch effects in the data. To minimize this, we used the integrate function in Seurat version 3.0 to cluster all cells from all samples together with 11dpa as the anchor data set with dims = 1:20 and all other parameters set to default (Butler et al., 2018; Stuart et al., 2019). The integrated dataset was then scaled and 30 principal components used for clustering with a resolution of 0.6 and visualized with tSNE. For sub-clustering of fibroblast and bone populations, fibroblast and bone clusters were subsetted from the integrated data set as Seurat objects and re-normalized. These objects were re-integrated in Seurat, again using 11dpa as the anchor data set and dims = 1:20, scaled, and clustered with principal components 1:20 and resolution 0.6 to reveal any subpopulations.

### Differential proportion analysis

Differential proportion analysis (Farbehi et al., 2019) was performed in R to statistically test for significant cluster membership over regenerative time. Cluster membership tables were calculated in Seurat and the resulting table of cells in each cluster by time point was used in differential proportion analysis. In the first step, generateNull was used with n = 100,000 and p = 0.1 as in the original reference. Significance values were calculated for pairwise comparisons of each time point with every other time point and were corrected for multiple hypothesis testing with the Benjamini-Hochberg method in R (Benjamini and Hochberg, 1995). Significance values reported in figures are: p < 0.05 (*), p < 0.01 (**), and p < 0.001 (***).

### Cell trajectory analyses

The SPRING web interface (https://kleintools.hms.harvard.edu/tools/spring.html) (Weinreb et al., 2018) was used to generate reproducible, continuous k nearest neighbors force-directed graphs of cells in gene expression space. A gene by cell expression matrix, a file containing time point and Seurat cluster metadata for each cell, and a list of gene names was the input to the web interface. All parameters were left at default values. Blastema datasets (11dpa, 12dpa, 14dpa, and 17dpa) were projected onto the unamputated dataset to avoid batch effects. Qualitative analysis of trajectories was facilitated by overlaying Seurat cluster information, regenerative stage, or gene expression. Differential gene expression associated with lineage trajectory (Figure 3, 4, and Supplemental Figure 7) was assessed in SPRING. Only genes with Z-score >1.96 were analyzed.

### Section RNA in situ hybridization

Adult wild-type CD1(ICR) (Charles River Laboratories) mice were used for all RNA in situ experiments. Blastema stage regenerating mouse digit tips, and contralateral unamputated controls, were collected and fixed in 4% paraformaldehyde at 4°C overnight, followed by washing and decalcification in decalcifying solution lite (Sigma Aldrich) (40 minutes at room temperature). Digits were prepared for embedding with a 5% to 30% sucrose gradient over 3 days, embedded in OCT (Tissue-Tek), and sectioned at 20µm on a Leica CM3050S cryostat. E12.5 embryos used for probe controls were collected from CD1(ICR) timed pregnant females, followed by PFA fixation and sucrose/OCT embedding as above, with solution change times of 30 minutes. A *Mest* cDNA for in situ probe template was PCR amplified from E10.5 mouse limb bud random-primed cDNA library with primers 5’GCTCCAGAACCGCAGAATCA and 5’GGGAGGTAATACAGGGAGGC (Mesman et al., 2018). The cDNA was cloned into the pGEM-T easy vector (Promega) and sequenced to confirm identity. Antisense RNA probe and sense negative control probe were generated by SP6 or T7 in vitro transcription with digoxigenin-UTP (Sigma Aldrich). Section RNA in situ hybridization was performed as previously reported (Murtaugh et al., 1999), with proteinase K used at 3ug/mL (room temp, 10 minutes). All digit tip in situ hybridized sections were developed for the same amount of time.

### Data and code availability

All single cell RNAseq FASTQ files and cell by gene expression matrices from this project are available in the NCBI Gene Expression Omnibus, Dataset Accession GSExxxxxx [upon publication]. No new computational tools were developed in this project, however the code for the usage of existing tools, as detailed above, is available at the permanent link: https://github.com/LehoczkyBWH/xxxxxx [upon publication].

## ACKNOWLEDGEMENTS

We appreciate helpful conversations with Drs. Michael Brenner and Soumya Raychaudhuri. We thank Drs. Caleb Weinreb and Allon Klein for help with SPRING data input. We are grateful for constructive comments from all members of the Lehoczky Lab. This work was supported by the Eunice Kennedy Shriver National Institute of Child and Human Development (R03HD093922 and R21HD097405 to J.A.L), The Osher Center for Integrative Medicine, and funds from BWH Department of Orthopedic Surgery to J.A.L.

## FIGURE LEGENDS AND TABLES

**Supplemental Figure 1.**
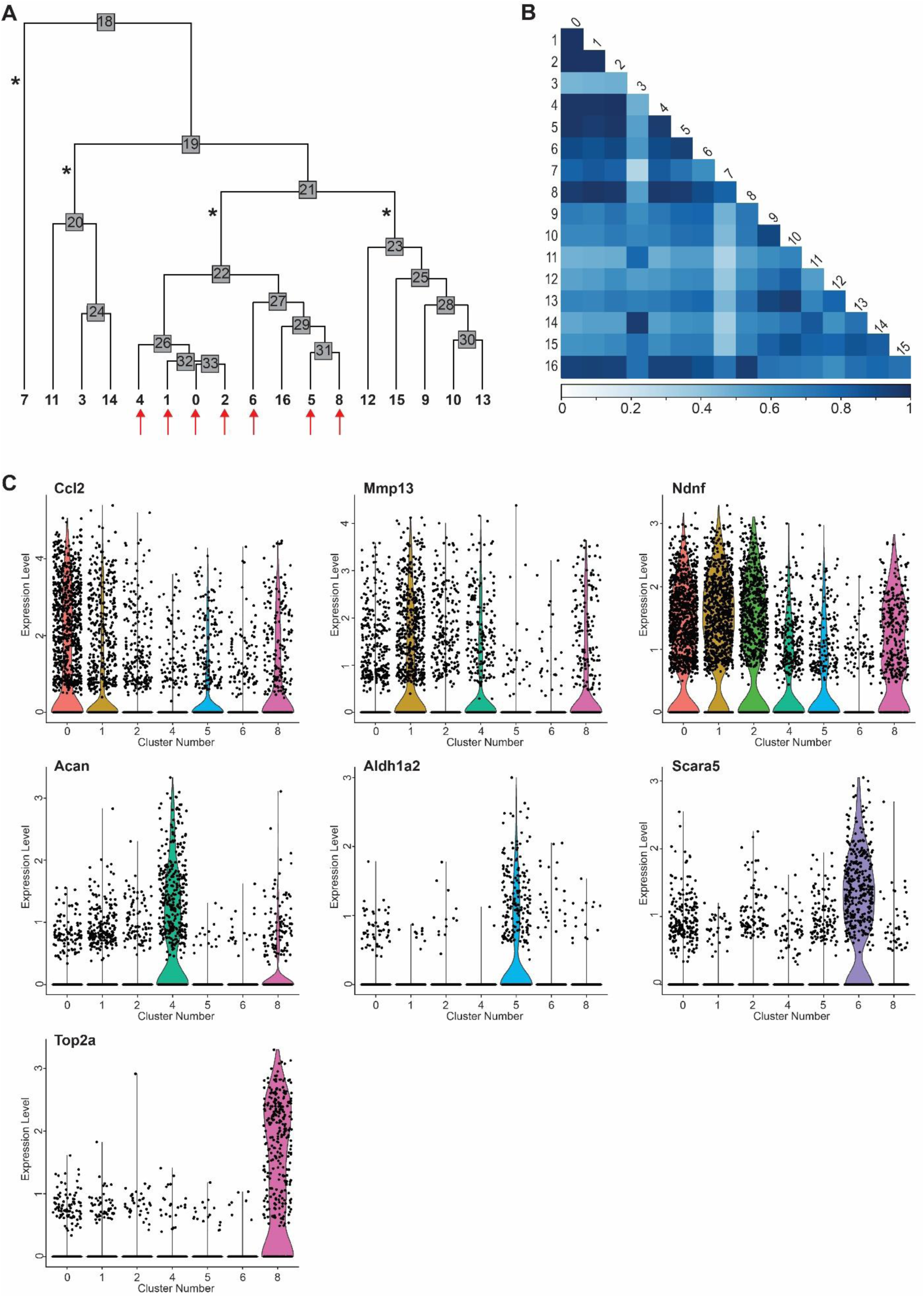
11dpa cluster relationships and fibroblast heterogeneity. Additional analyses of data presented in Figure 1. (A) Dendrogram showing the relationships between clusters; cell cluster numbers correlate with tSNE plot cluster numbers in Figure 1B. Asterisks denote four main branches of the dendrogram, and red arrows denote fibroblast clusters. (B) Heatmap showing the Pearson correlation between each cell cluster, where dark blue represents highly correlated (r nears 1) and light blue represents lowly correlated (r nears 0). (C) Violin plots of representative genes differentially expressed among fibroblast clusters. Black points represent individual cells and the colored curve shows the distribution of cells at a given expression level. Examples include: *Ccl2* (cluster 0), *Mmp13* (cluster 1), *Ndnf* (clusters 0, 1, 2, and 8), *Acan* (cluster 4), *Aldh1a2* (cluster 5), *Scara5* (cluster 6), and *Top2a* (cluster 8).

**Supplemental Figure 2.**
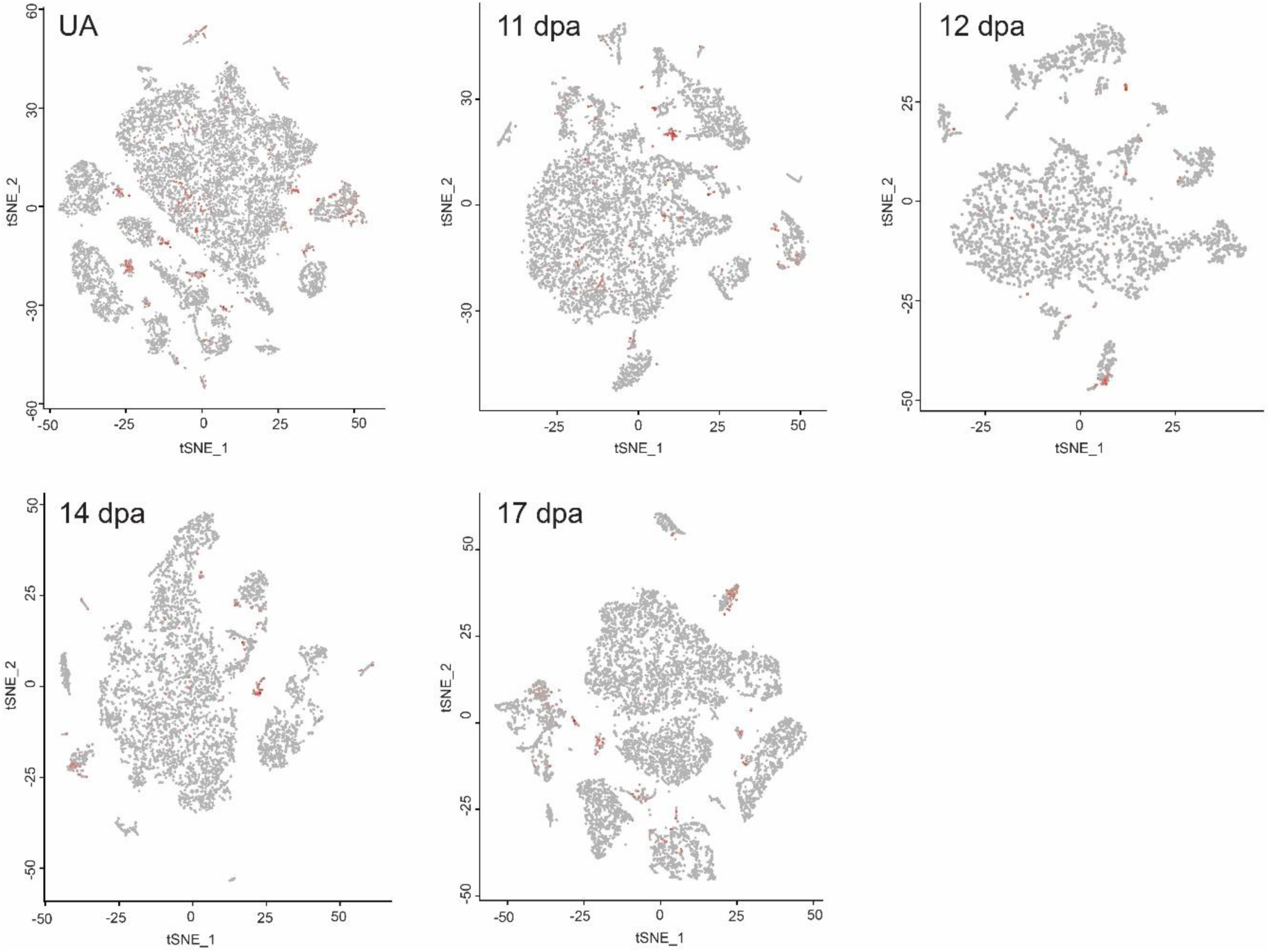
Visualization of predicted doublet cells at each regenerative time point. tSNE plots for each blastema regenerative timepoint: unamputated (UA) (corresponds to data in Supplemental Figure 6A), 11dpa (corresponds to data in Figure 1A), 12dpa (corresponds to data in Supplemental Figure 3A), 14dpa (corresponds to data in Supplemental Figure 4A), and 17 dpa (corresponds to data in Supplemental Figure 5A). 419 cells were classified as doublets in the 11dpa sample, 83 in 12dpa, 271 in 14dpa, 606 in 17dpa, and 978 in UA. Cells classified as doublets and excluded from all subsequent analyses are in red; all other cells are in gray.

**Supplemental Figure 3.**
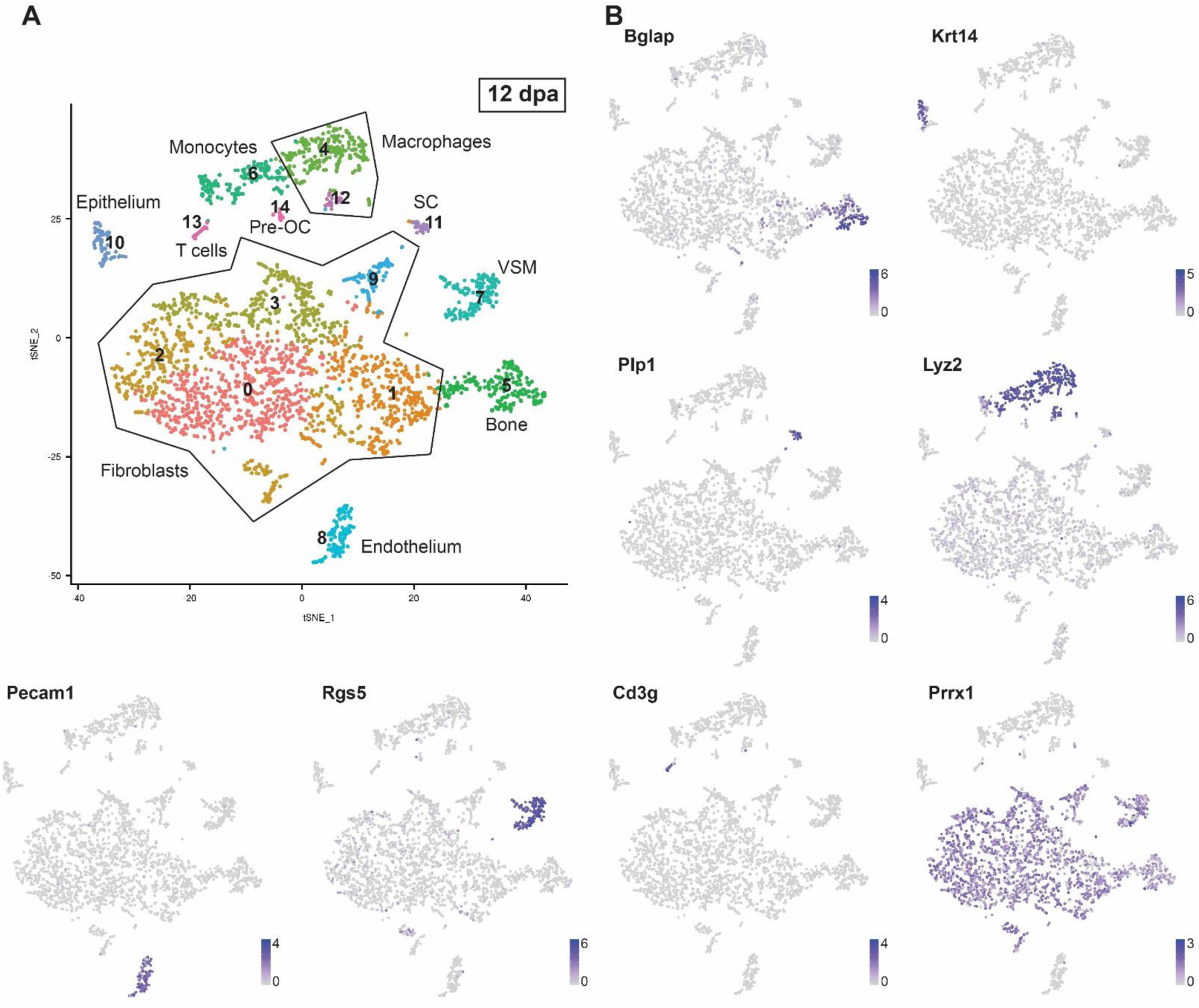
Single cell RNAseq of 12dpa blastema. (A) Unbiased single cell clustering of 3,309 high quality cells visualized by tSNE plot. Each dot represents a single cell and cells assigned to the same cluster are similarly colored. Cell type identities are assigned as follows: fibroblasts (clusters 0-3, and 9), macrophages (clusters 4 and 12), bone (cluster 5), monocytes (cluster 6), vascular smooth muscle cells (VSM) (cluster 7), endothelial cells (cluster 8), epithelial cells (cluster 10), Schwann cells (SC) (cluster 11), T cells (cluster 13), and pre-osteoclasts (Pre-OC) (cluster 14). (B) Gene expression tSNE overlay with examples of highly expressed, cell type specific markers used to assign cluster cell identities: *Bglap* (bone), *Krt14* (epithelial cells), *Plp1* (SCs), *Lyz2* (macrophages), *Pecam1* (endothelial cells), *Rgs5* (vascular smooth muscle cells), *Cd3g* (T cells), *Prrx1* (fibroblasts). Gray depicts low expression and purple depicts high expression as specified on the scale for each gene.

**Supplemental Figure 4.**
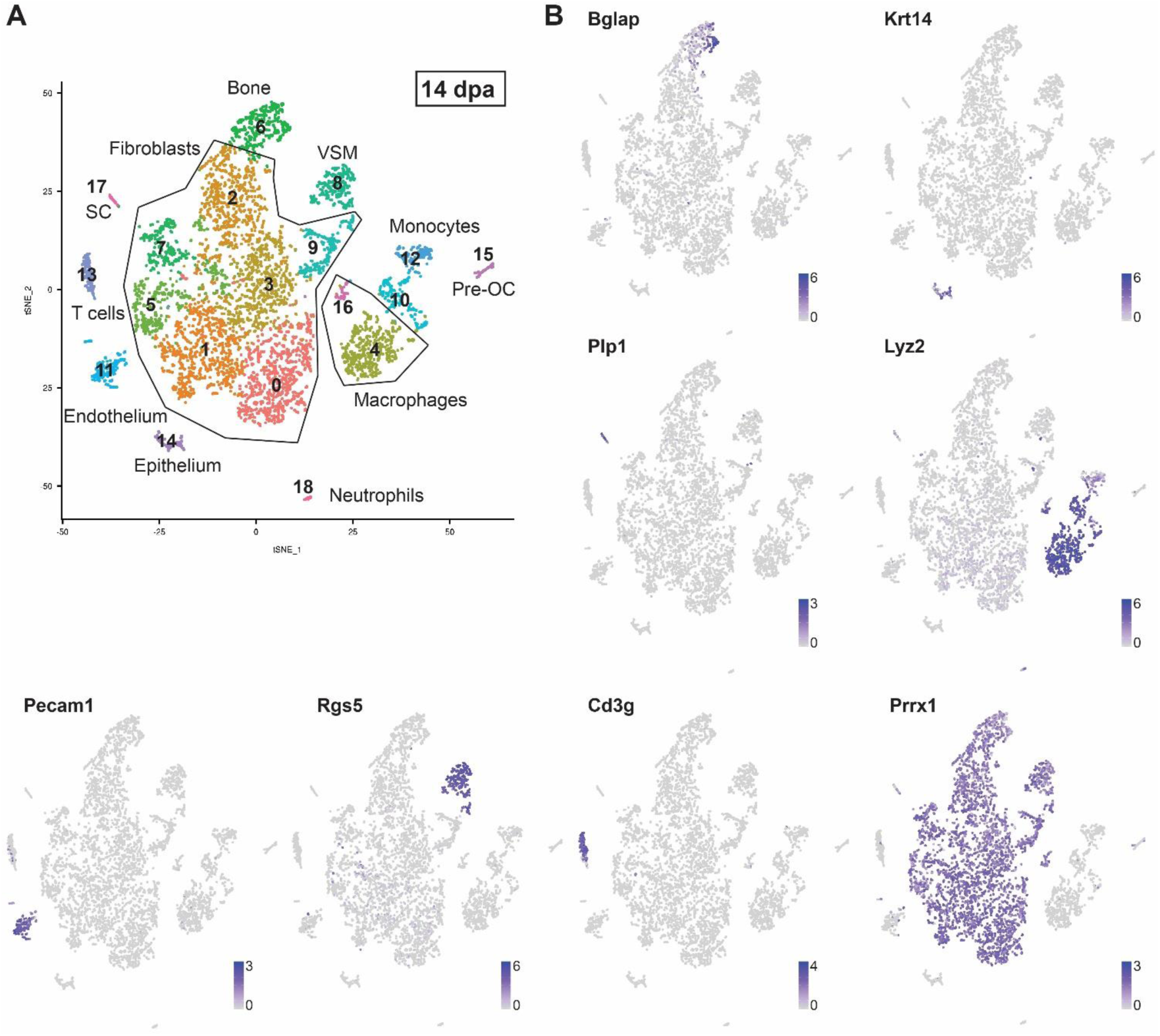
Single cell RNAseq of 14dpa blastema. (A) Unbiased single cell clustering of 5,896 high quality cells visualized by tSNE plot. Each dot represents a single cell and cells assigned to the same cluster are similarly colored. Cell type identities are assigned as follows: fibroblasts (clusters 0-3, 5, 7, and 9), macrophages (clusters 4 and 16), bone (cluster 6), vascular smooth muscle cells (VSM) (cluster 8), monocytes (clusters 10 and 12), endothelial cells (cluster 11), T cells (cluster 13), epithelial cells (cluster 14), pre-osteoclast (Pre-OC) (cluster 15), Schwann cells (SC) (cluster 17), neutrophils (cluster 18). (B) Gene expression tSNE overlay with examples of highly expressed, cell type specific markers used to assign cluster cell identities: *Bglap* (bone), *Krt14* (epithelial cells), *Plp1* (SCs), *Lyz2* (macrophages), *Pecam1* (endothelial cells), *Rgs5* (vascular smooth muscle cells), *Cd3g* (T cells), *Prrx1* (fibroblasts). Gray depicts low expression and purple depicts high expression as specified on the scale for each gene.

**Supplemental Figure 5.**
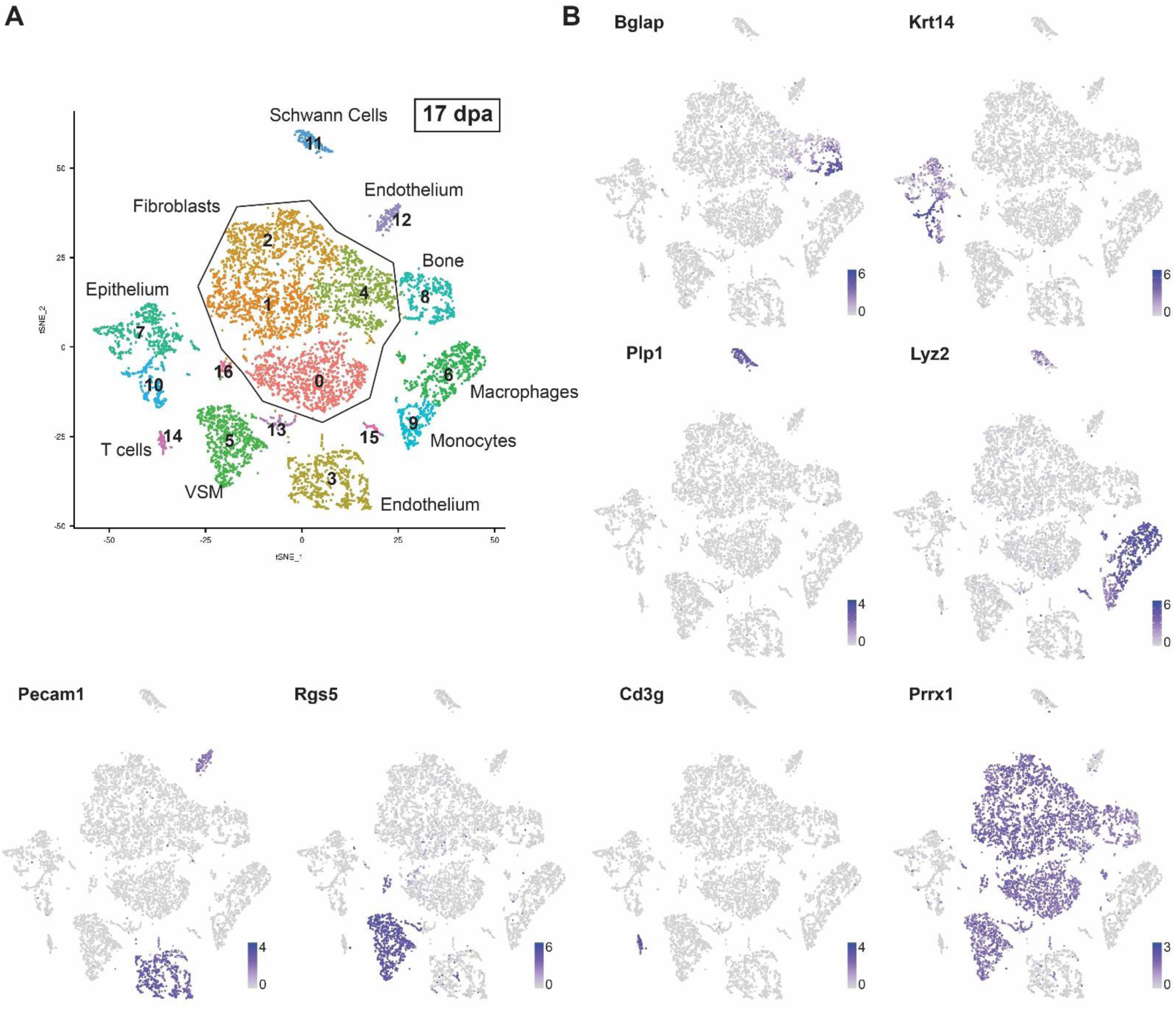
Single cell RNAseq of 17dpa blastema. (A) Unbiased single cell clustering of 8,778 high quality cells visualized by tSNE plot. Each dot represents a single cell and cells assigned to the same cluster are similarly colored. Cell type identities are assigned as follows: fibroblasts (clusters 0-2, and 4), endothelial cells (cluster 3), vascular smooth muscle cells (VSM) (clusters 5, 13 and 16), macrophages (cluster 6), epithelial cells (clusters 7 and 10), bone (cluster 8), monocytes (clusters 9 and 15), Schwann cells (SC) (cluster 11), endothelial cells (cluster 12), T cells (cluster 14). (B) Gene expression tSNE overlay with examples of highly expressed, cell type specific markers used to assign cluster cell identities: *Bglap* (bone), *Krt14* (epithelial cells), *Plp1* (SCs), *Lyz2* (macrophages), *Pecam1* (endothelial cells), *Rgs5* (vascular smooth muscle cells), *Cd3g* (T cells), *Prrx1* (fibroblasts). Gray depicts low expression and purple depicts high expression as specified on the scale for each gene.

**Supplemental Figure 6.**
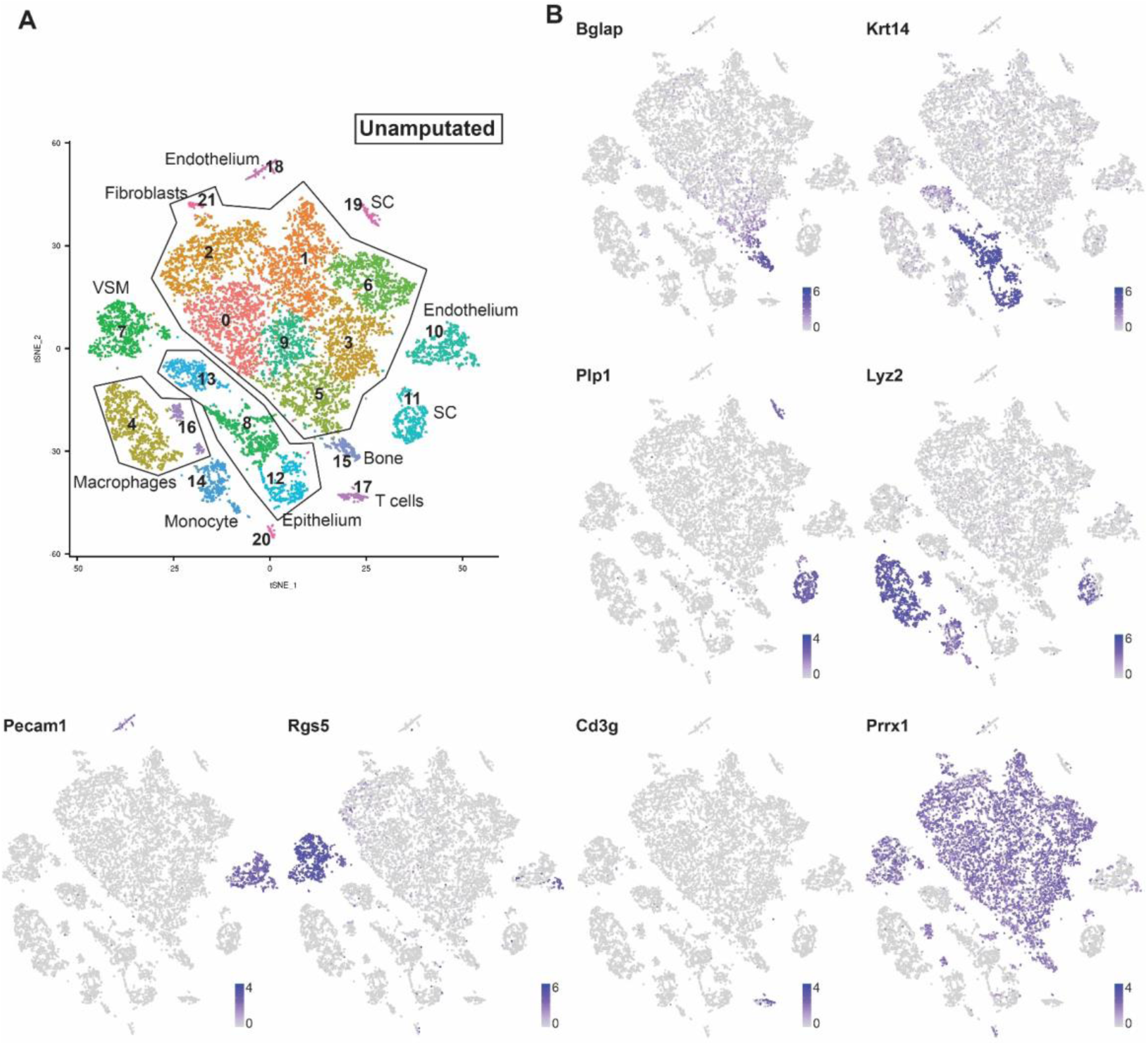
Single cell RNAseq of the unamputated digit tip. (A) Unbiased single cell clustering of 12,871 high quality cells visualized by tSNE plot. Each dot represents a single cell and cells assigned to the same cluster are similarly colored. Cell type identities are assigned as follows: fibroblasts (clusters 0-3, 5, 6, 9, 20 and 21), macrophages (clusters 4 and 16), vascular smooth muscle cells (VSM) (cluster 7), epithelial cells (clusters 8, 12, and 13), endothelial cells (cluster 10 and 18), Schwann cells (SC) (clusters 11 and 19), monocytes (cluster 14), bone (cluster 15), T cells (cluster 17). (B) Gene expression tSNE overlay with examples of highly expressed, cell type specific markers used to assign cluster cell identities: *Bglap* (bone), *Krt14* (epithelial cells), *Plp1* (SCs), *Lyz2* (macrophages), *Pecam1* (endothelial cells), *Rgs5* (vascular smooth muscle cells), *Cd3g* (T cells), *Prrx1* (fibroblasts). Gray depicts low expression and purple depicts high expression as specified on the scale for each gene.

**Supplemental Figure 7.**
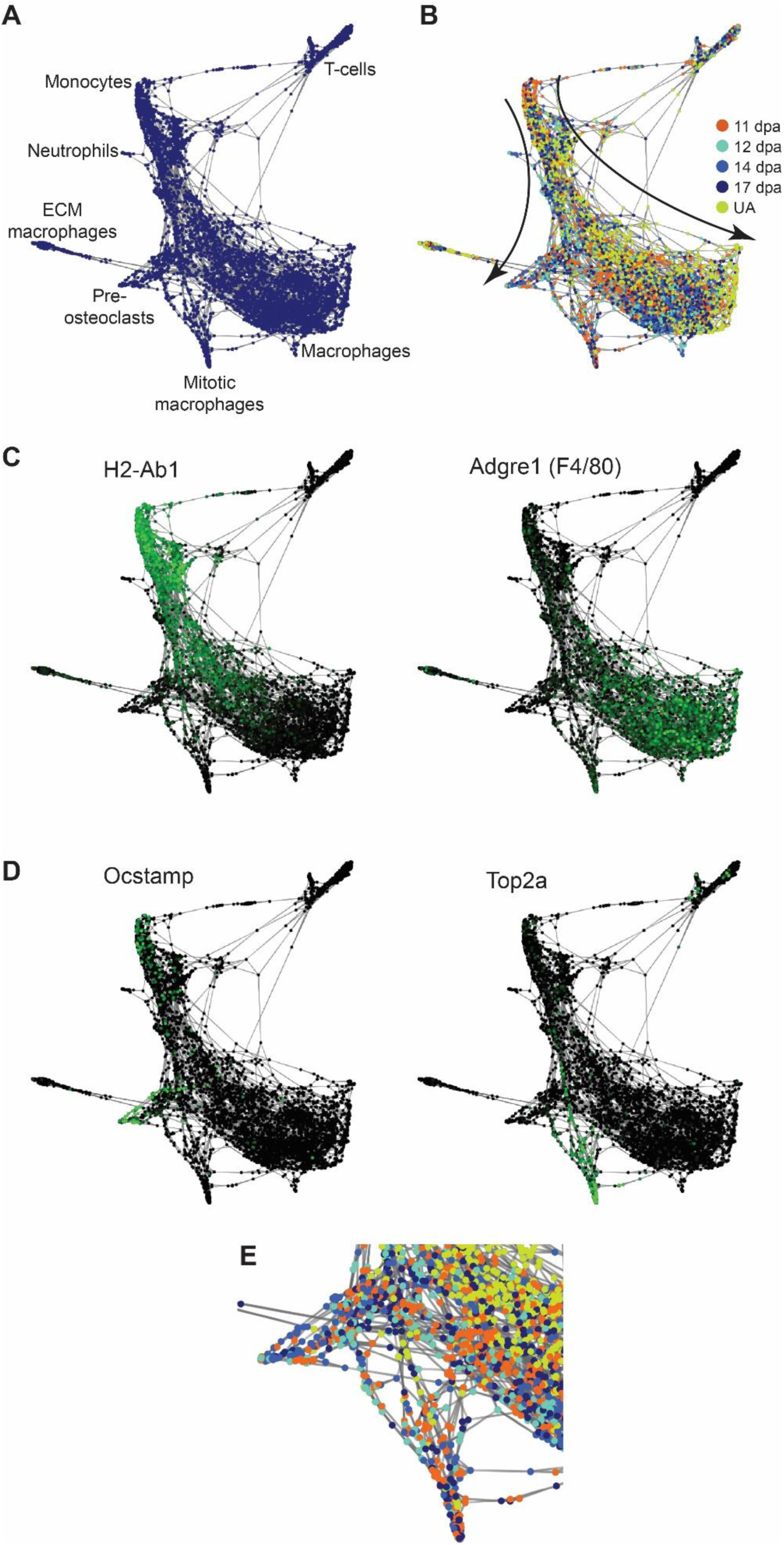
Differentiation trajectory analysis of immune-related cells. SPRING lineage trajectory analysis of cells from the integrated data set immune-related clusters 7, 11, 12, 17, 18, 21, and 23. (A) Force-directed plot of cells showing a monocytes, macrophages, ECM macrophages (express ECM related genes; population of unknown relevance), mitotic macrophages, pre-osteoclasts, T cells. (B) SPRING plot as in (A) with regenerative stages of each cell colored coded: 11dpa (orange), 12dpa (light blue), 14dpa (medium blue), 17dpa (dark blue), unamputated (yellow). Known differentiation trajectories from monocytes to macrophages, and monocytes to pre-osteoclasts are depicted with curved arrows. (C) Gene expression overlay showing monocyte to macrophage differentiation. *H2-Ab1* is expressed in monocytes and *Adgre1*(F4/80) is expressed in macrophages. High expression is in green and low expression is black. (D) Gene expression overlay showing *Ocstamp* pre-osteoclast expression and *Top2a* mitotic macrophage expression. (E) Close-up of pre-osteoclasts and mitotic macrophages in (B); qualitative evaluation shows enrichment for blastema stages.

**Supplemental Figure 8.**
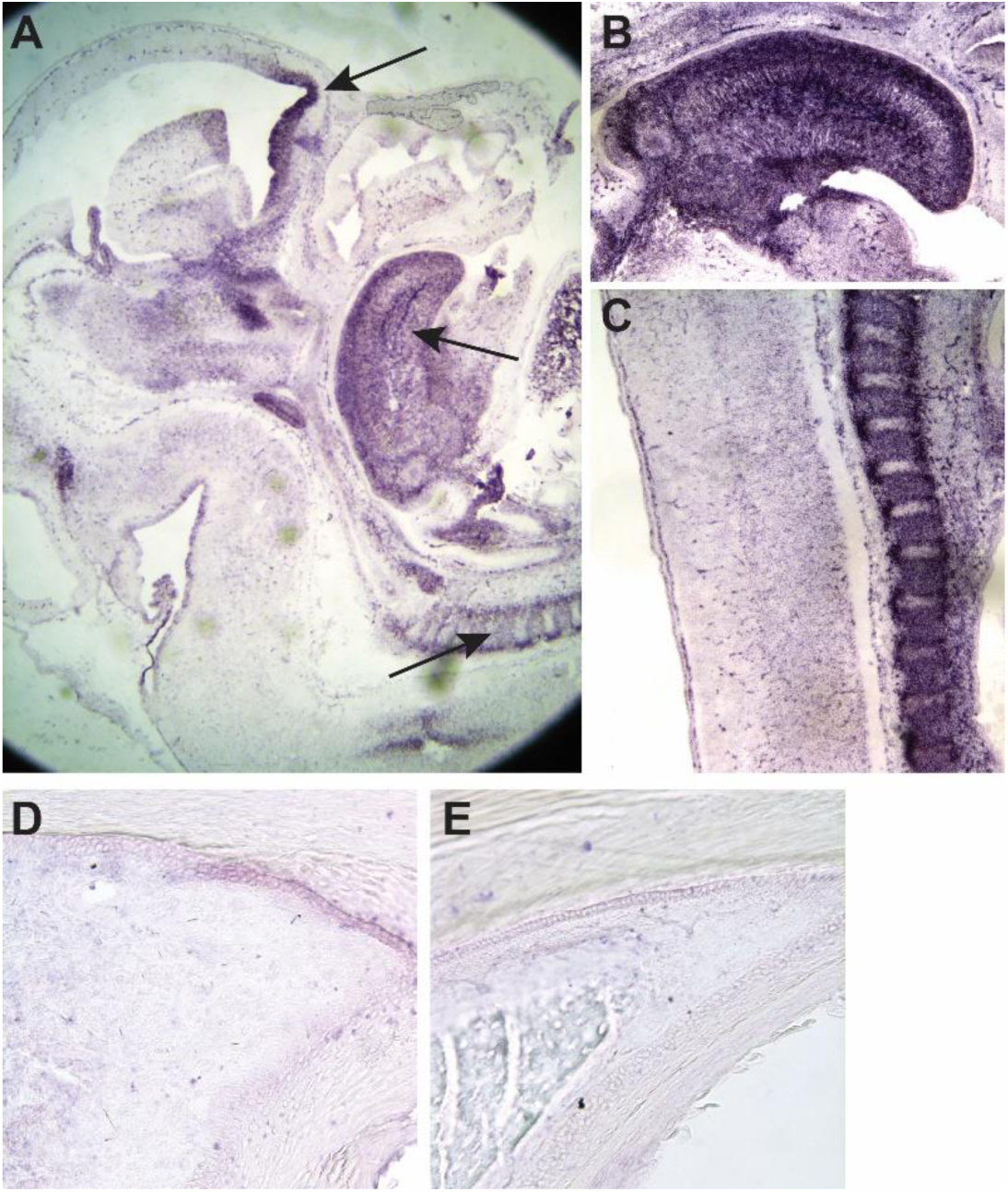
*Mest* RNA in situ control expression. All panels are DIG labeled section RNA in situ controls. (A-C) *Mest* antisense probe positive control on E12.5 mouse embryo sections. (A) Transverse section through head and neck region with positive expression (purple) in the developing forebrain, tongue, and vertebrae (arrows). Magnified view of panel (A) of (B) tongue and (C) vertebrae. (D and E) *Mest* sense probe negative control on adult mouse digit tip sections. No appreciable expression is found on (D) 12dpa or (E) unamputated tissues.

**Supplemental Table 1**

**Cell cluster differential gene expression by regenerative stage**

Differential gene expression analysis output from the FindAllMarkers function in Seurat. Each tab contains data from discrete regenerative stages: (A) 11dpa, (B) 12 dpa, (C) 14dpa, (D) 17dpa, and (E) unamputated. Column headers are: gene (NCBI gene ID), p-val (unadjusted p-value), avg logFC (average log fold-change among all cell clusters at this stage), pct.1 (percentage of cells in this cluster with this gene expression), pct.2 (percentage of cells in all other clusters with this gene expression), adj p-val (Bonferroni corrected p-value), cluster (cell cluster number on associated tSNE plot), cell type (cell cluster associated cell type assigned by literature review of most significant genes).

**Supplemental Table 2.**
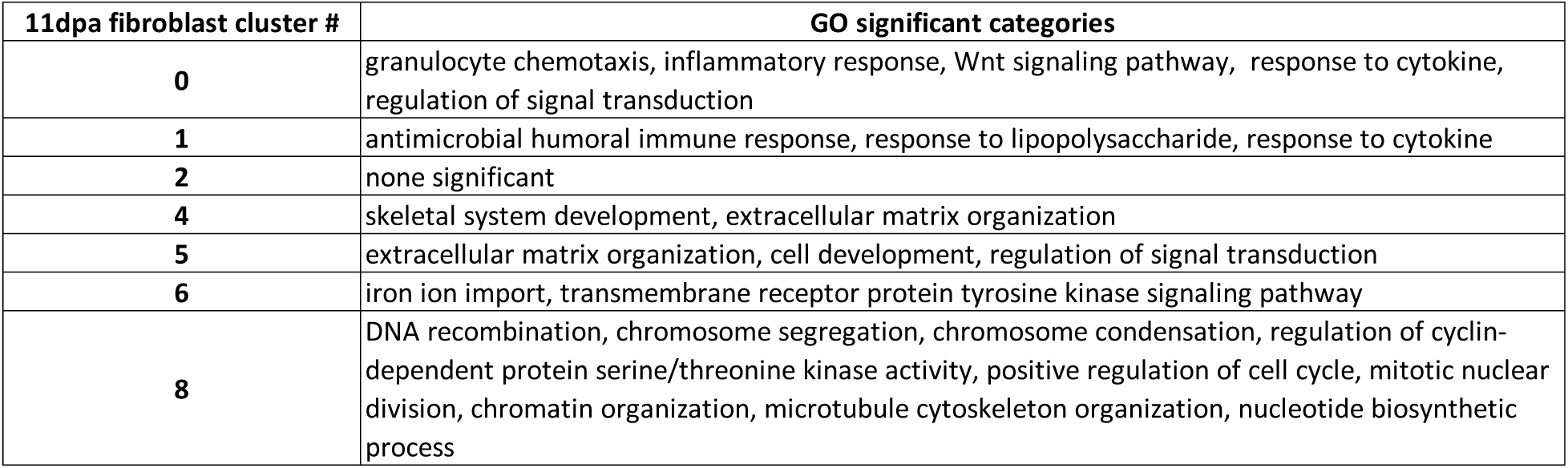
GO terms associated with 11dpa fibroblast cluster gene expression. Significant GO slim biological process categories for 11dpa fibroblast clusters with adjusted p- value ≤ 0.05 and average log fold-change ≥ 0.05.

| 11dpa fibroblast cluster # | GO significant categories |
| --- | --- |
| 0 | granulocyte chemotaxis, inflammatory response, Wnt signaling pathway, response to cytokine, regulation of signal transduction |
| 1 | antimicrobial humoral immune response, response to lipopolysaccharide, response to cytokine |
| 2 | none significant |
| 4 | skeletal system development, extracellular matrix organization |
| 5 | extracellular matrix organization, cell development, regulation of signal transduction |
| 6 | iron ion import, transmembrane receptor protein tyrosine kinase signaling pathway |
| 8 | DNA recombination, chromosome segregation, chromosome condensation, regulation of cyclin-dependent protein serine/threonine kinase activity, positive regulation of cell cycle, mitotic nuclear division, chromatin organization, microtubule cytoskeleton organization, nucleotide biosynthetic process |

**Supplemental Table 3**

**Integrated data set cell cluster differential gene expression**

Differential gene expression analysis of 11dpa, 12dpa, 14dpa, 17dpa, and unamputated integrated data set. Output is from the FindAllMarkers function in Seurat. Column headers are: gene (NCBI gene ID), p-val (unadjusted p-value), avg logFC (average log fold-change among all cell clusters at this stage), pct.1 (percentage of cells in this cluster with this gene expression), pct.2 (percentage of cells in all other clusters with this gene expression), adj p-val (Bonferroni corrected p-value), cluster (cell cluster number on associated tSNE plot), cell type (cell cluster associated cell type assigned by literature review of most significant genes).

**Supplemental Table 4.** P-values for differential population analysis. All resultant p-values for regenerative stage pairwise differential proportion analyses testing for significant changes in proportion of cells within clusters. Reported values have been corrected for multiple hypothesis testing. Column headers indicate regenerative stages being compared: 11 = 11dpa, 12 = 12dpa, 14 = 14dpa, 17 = 17dpa, and ua = unamputated. Cluster numbers in each row refer to tSNE cluster classification in Figure 2B. All table cells in gray are noted as significant with p ≤ 0.05.

|  | ua vs. 11 | ua vs. 12 | ua vs. 14 | ua vs. 17 | 11 vs. 12 | 11 vs. 14 | 11 vs. 17 | 12 vs. 14 | 12 vs. 17 | 14 vs. 17 |
| --- | --- | --- | --- | --- | --- | --- | --- | --- | --- | --- |
| cluster_0 | 0.0023 | 0.0568 | 0.0568 | 0.4455 | 0.3148 | 0.1976 | 0.0023 | 0.4455 | 0.0568 | 0.0568 |
| cluster_1 | 0.0000 | 0.0010 | 0.0002 | 0.1390 | 0.1390 | 0.0746 | 0.0000 | 0.4276 | 0.0124 | 0.0059 |
| cluster_2 | 0.1002 | 0.2819 | 0.1362 | 0.1645 | 0.4646 | 0.4725 | 0.4646 | 0.4646 | 0.4973 | 0.4646 |
| cluster_3 | 0.2321 | 0.2321 | 0.2742 | 0.2742 | 0.3901 | 0.4293 | 0.4258 | 0.3454 | 0.3425 | 0.4646 |
| cluster_4 | 0.0028 | 0.1201 | 0.1296 | 0.1296 | 0.3100 | 0.1296 | 0.1201 | 0.3317 | 0.3117 | 0.4456 |
| cluster_5 | 0.4473 | 0.4719 | 0.4719 | 0.4719 | 0.4719 | 0.4473 | 0.4473 | 0.4719 | 0.4719 | 0.4719 |
| cluster_6 | 0.1108 | 0.2819 | 0.1604 | 0.0384 | 0.4438 | 0.4438 | 0.0029 | 0.4438 | 0.0289 | 0.0060 |
| cluster_7 | 0.4243 | 0.4243 | 0.4243 | 0.4243 | 0.4256 | 0.4243 | 0.4243 | 0.4243 | 0.4243 | 0.4243 |
| cluster_8 | 0.1539 | 0.1363 | 0.0815 | 0.1539 | 0.2702 | 0.2702 | 0.4995 | 0.4995 | 0.2702 | 0.2702 |
| cluster_9 | 0.4636 | 0.4947 | 0.2743 | 0.0002 | 0.4636 | 0.2688 | 0.0011 | 0.3520 | 0.0031 | 0.0000 |
| cluster_10 | 0.0005 | 0.0015 | 0.0001 | 0.0631 | 0.3808 | 0.2893 | 0.0767 | 0.3808 | 0.0767 | 0.0270 |
| cluster_11 | 0.4779 | 0.4779 | 0.4779 | 0.4779 | 0.4779 | 0.4779 | 0.4779 | 0.4779 | 0.4779 | 0.4779 |
| cluster_12 | 0.4540 | 0.4540 | 0.4540 | 0.4540 | 0.4540 | 0.4540 | 0.4540 | 0.4540 | 0.4540 | 0.4540 |
| cluster_13 | 0.0000 | 0.0100 | 0.0086 | 0.3184 | 0.2567 | 0.0846 | 0.0001 | 0.3184 | 0.0192 | 0.0192 |
| cluster_14 | 0.0285 | 0.0853 | 0.0066 | 0.1366 | 0.4755 | 0.3185 | 0.3125 | 0.3185 | 0.3185 | 0.1366 |
| cluster_15 | 0.0095 | 0.1772 | 0.0095 | 0.3703 | 0.2934 | 0.4438 | 0.0359 | 0.2910 | 0.2850 | 0.0359 |
| cluster_16 | 0.0000 | 0.0038 | 0.0003 | 0.0652 | 0.4495 | 0.4735 | 0.0505 | 0.4495 | 0.1109 | 0.0568 |
| cluster_17 | 0.4860 | 0.4860 | 0.1714 | 0.4860 | 0.4860 | 0.1714 | 0.4860 | 0.1879 | 0.4860 | 0.1891 |
| cluster_18 | 0.4443 | 0.4443 | 0.4443 | 0.4443 | 0.4756 | 0.4443 | 0.4443 | 0.4443 | 0.4443 | 0.4443 |
| cluster_19 | 0.3982 | 0.4135 | 0.3982 | 0.3982 | 0.3982 | 0.4135 | 0.3892 | 0.4135 | 0.3982 | 0.3982 |
| cluster_20 | 0.0460 | 0.0460 | 0.0460 | 0.0460 | 0.4745 | 0.4745 | 0.4745 | 0.4745 | 0.4745 | 0.4745 |
| cluster_21 | 0.2324 | 0.2324 | 0.1822 | 0.2997 | 0.3372 | 0.2745 | 0.2997 | 0.3637 | 0.2745 | 0.2324 |
| cluster_22 | 0.3231 | 0.3231 | 0.3231 | 0.3231 | 0.3231 | 0.3231 | 0.4699 | 0.4699 | 0.3231 | 0.3231 |
| cluster_23 | 0.3733 | 0.3733 | 0.3920 | 0.3920 | 0.3920 | 0.3733 | 0.3733 | 0.3733 | 0.3733 | 0.3733 |

**Supplemental Table 5**

**Differential gene expression from all-stage integrated and re-clustered fibroblasts and bone**

Differential gene expression analysis from only clustering of only fibroblast and bone cells of 11dpa, 12dpa, 14dpa, 17dpa, and unamputated digit tip data. Output is from the FindAllMarkers function in Seurat. Column headers are: gene (NCBI gene ID), p-val (unadjusted p-value), avg logFC (average log fold-change among all cell clusters at this stage), pct.1 (percentage of cells in this cluster with this gene expression), pct.2 (percentage of cells in all other clusters with this gene expression), adj p-val (Bonferroni corrected p-value), cluster (cell cluster number on associated tSNE plot (Figure 4A)), cell type (cell cluster associated cell type assigned by literature review of most significant genes).

**Supplemental Table 6.** P-values for differential population analysis of re-clustered fibroblast and bone populations. All resultant p-values for regenerative stage pairwise differential proportion analyses testing for significant changes in proportion of cells within clusters. Reported values have been corrected for multiple hypothesis testing. Column headers indicate regenerative stages being compared: 11 = 11dpa, 12 = 12dpa, 14 = 14dpa, 17 = 17dpa, and ua = unamputated. Cluster numbers in each row refer to tSNE cluster classification in Figure 4A. All table cells in gray are noted as significant with p ≤ 0.05.

|  | ua vs. 11 | ua vs. 12 | ua vs. 14 | ua vs. 17 | 11 vs. 12 | 11 vs. 14 | 11 vs. 17 | 12 vs. 14 | 12 vs. 17 | 14 vs. 17 |
| --- | --- | --- | --- | --- | --- | --- | --- | --- | --- | --- |
| cluster_0 | 0.0000 | 0.0009 | 0.0006 | 0.0405 | 0.1726 | 0.0554 | 0.0006 | 0.3089 | 0.0554 | 0.0785 |
| cluster_1 | 0.1733 | 0.4456 | 0.4456 | 0.4456 | 0.3454 | 0.2305 | 0.2156 | 0.4456 | 0.4456 | 0.4456 |
| cluster_2 | 0.0104 | 0.0812 | 0.0359 | 0.3688 | 0.3688 | 0.3913 | 0.0409 | 0.4161 | 0.1740 | 0.0812 |
| cluster_3 | 0.4408 | 0.4408 | 0.4408 | 0.4408 | 0.4408 | 0.4408 | 0.4408 | 0.4408 | 0.4408 | 0.4408 |
| cluster_4 | 0.0015 | 0.0345 | 0.0722 | 0.2622 | 0.3607 | 0.1418 | 0.0188 | 0.2622 | 0.0931 | 0.1814 |
| cluster_5 | 0.2015 | 0.0477 | 0.0825 | 0.3363 | 0.1497 | 0.2620 | 0.2828 | 0.2620 | 0.0825 | 0.1591 |
| cluster_6 | 0.0004 | 0.0052 | 0.0081 | 0.0677 | 0.4803 | 0.2895 | 0.0911 | 0.3230 | 0.1509 | 0.2734 |
| cluster_7 | 0.4862 | 0.4962 | 0.4862 | 0.4862 | 0.4862 | 0.4862 | 0.4862 | 0.4862 | 0.4862 | 0.4962 |
| cluster_8 | 0.3203 | 0.1738 | 0.1738 | 0.1738 | 0.3203 | 0.3203 | 0.3203 | 0.4540 | 0.4540 | 0.4890 |
| cluster_9 | 0.4461 | 0.4461 | 0.4461 | 0.4461 | 0.4461 | 0.4461 | 0.4461 | 0.4461 | 0.4461 | 0.4461 |
| cluster_10 | 0.4364 | 0.0568 | 0.0198 | 0.0288 | 0.0674 | 0.0219 | 0.0414 | 0.4364 | 0.4364 | 0.3863 |
| cluster_11 | 0.0189 | 0.0366 | 0.0366 | 0.4015 | 0.4406 | 0.4015 | 0.0366 | 0.4015 | 0.0603 | 0.0620 |
| cluster_12 | 0.0000 | 0.0001 | 0.0000 | 0.0018 | 0.4978 | 0.4978 | 0.2252 | 0.4978 | 0.2252 | 0.2252 |
| cluster_13 | 0.0116 | 0.3093 | 0.2400 | 0.3513 | 0.1733 | 0.1588 | 0.0489 | 0.4763 | 0.3703 | 0.3513 |
| cluster_14 | 0.0996 | 0.0996 | 0.0996 | 0.0996 | 0.4437 | 0.4437 | 0.4437 | 0.4437 | 0.4437 | 0.4437 |

**Supplemental Table 7**

**Differential gene expression of blastema-enriched versus blastema-depleted cell clusters**

Differential gene expression analysis from defined blastema-enriched clusters (Figure 4C) versus blastema-depleted clusters (Figure 4B). Output is from the FindAllMarkers function in Seurat. Column headers are: gene (NCBI gene ID), p-val (unadjusted p-value), avg logFC (average log fold-change among all cell clusters at this stage), pct.1 (percentage of cells in blastema-enriched clusters with detected expression of the gene), pct.2 (percentage of cells in blastema-depleted clusters with detected expression of the gene), and adj p-val (Bonferroni corrected p-value).

**Supplemental Table 8.** Metadata for single cell RNAseq data thresholding. Summary of thresholding parameters for data processing, quality control, and cell clustering of each single cell RNAseq data set. Column headers denote blastema datasets (11dpa, 12dpa, 14dpa, and 17dpa) and unamputated control (UA). Parameters are: percent of mitochondrial genes upper bound, number of unique molecular identifiers (nUMI) lower and upper bounds, number of principal components, and resolution.

|  | 11dpa | 12dpa | 14dpa | 17dpa | UA |
| --- | --- | --- | --- | --- | --- |
| % Mitochondrial genes | 25 | 20 | 20 | 25 | 25 |
| nUMI lower bound | 200 | 200 | 200 | 200 | 200 |
| nUMI upper bound | 5500 | 4000 | 6500 | 6500 | 5000 |
| # of principal components | 16 | 16 | 20 | 20 | 20 |
| Resolution | 0.6 | 0.6 | 0.6 | 0.6 | 0.6 |

